# Metabolic Complementation between Glucose and Amino Acid Drives Hepatic *De Novo* Lipogenesis and Steatosis

**DOI:** 10.1101/2021.05.08.443229

**Authors:** Yilie Liao, Lei Liu, Honghao Li, Xiaojie Bai, Fangfang Sun, Xia Xiao, Suneng Fu

## Abstract

Increased *de novo* lipogenesis (DNL) is a hallmark of nonalcoholic fatty liver disease (NAFLD) in obesity, but the macronutrient source for >80% carbon backbone for fatty acid synthesis has not been determined. Here we take an integrated approach to dissect nutrient metabolism, both *ex vivo* and *in vivo*. We discover a castling effect of glucose and glutamine metabolism through *ex vivo* isotope tracing studies that limits the entrance of glucose carbon into the glutamine-dominated tricarboxylic acid cycle (TCA) and DNL pathways. *In vivo* tracing studies with a high carbohydrate drink (glucose/amino acid, 3:1, *w/w*) confirm dietary amino acids are twice more efficient than glucose in labeling the hepatic acetyl-CoA and fatty acid pool, and together they account for over 70% of hepatic DNL substrate. Both glucose and glutamine carbon flux into DNL pathways are increased in obese hepatocytes, and metabolic rerouting of substrate carbon toward glycogen synthesis and energy production through GYS2 and GLUD1 overexpression improves hepatic steatosis. Together, these data reveal the quantitative contribution of glucose and amino acid carbon toward hepatic DNL and the development of hepatic steatosis in obesity.

## Introduction

Nonalcoholic fatty liver disease (NAFLD) encompasses a continuum of liver pathology that starts with excessive triglyceride accumulation (hepatic steatosis) and gradually progresses toward inflammation (steatotic hepatitis), fibrosis, and even cirrhosis^1, 2^. NAFLD is strongly associated with diabetes^3^ and cardiovascular diseases^4^, and it is the most frequent cause of abnormal liver function, accounts for more than 25% of hepatic carcinoma^5^ and ∼20% of end-stage liver failure^6^. The global prevalence of NAFLD has reached 25%, but there is no drug presently on the market for treatment^7, 8^.

Obesity and insulin resistance are the leading causes of NAFLD^7^. Obesity may promote the development of NAFLD by supplying adipose-derived free fatty acids to the liver. It is estimated that more than 60% of fatty acids in the liver is sourced from the adipose tissue^9^, and adipose lipolysis is increased in obese and insulin resistant state^10, 11^. Re-esterification of adipose-derived free fatty acids into triglyceride requires glycerol moieties that may be sourced from the adipose or synthesized in situ through glyceroneogenesis and glycolytic pathways^12–14^. DNL is upregulated in animal models of obesity as well as obese patients, and it is the only source of fatty acid that is significantly increased under obese conditions^15^.

The obese state drives DNL through multiple mechanisms, but the principle substrate source of DNL and its regulation in the obese state remains unsettled^16^. Work by Clark and colleagues showed that glucose and fructose were not suitable for DNL in primary hepatocytes when compared to lactate, pyruvate, acetate, and alanine^17^. In rat intravenous infusion models, the estimated contribution of glucose toward cytosolic acetyl-CoA synthesis and lipogenesis ranged from 70% to 3∼5%^18–20^. In human, the majority of excess carbohydrate is stored as glycogen, and only 1-2% of a carbohydrate meal is converted to lipids^21, 22^. A recent fluxomics study of 15 nutrients fail to label fatty acids *in vivo*^23^.

Through a series of flux and genetic analyses, here we examine the quantitative contribution of glucose and amino acids toward fatty acid synthesis and the development of hepatic steatosis in obesity.

### Hepatic glucose flux is restricted from the TCA cycle

Glucose has many metabolic fates and it is remotely placed from the tricarboxylic acid (TCA) cycle and the *de novo* lipogenesis (DNL) pathway (Fig. 1a). We performed flux analysis of uniformly-labeled glucose ([U-^13^C]Glc) to track its kinetics and fate of metabolism in primary hepatocytes and examine its potential remodeling during the development of nonalcoholic fatty liver disease (NAFLD) in the leptin-deficient, *ob/ob* mice model.

**Figure 1.**
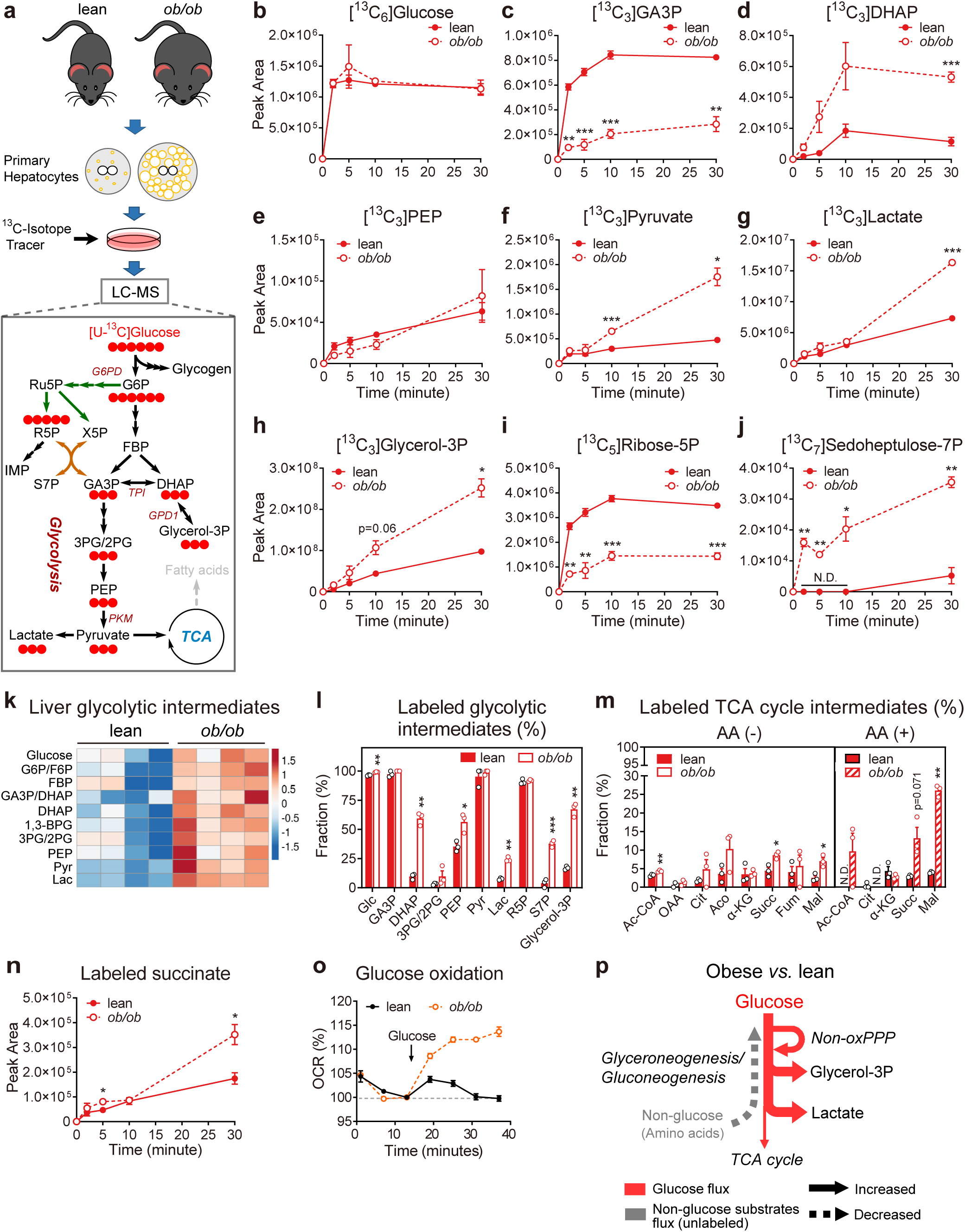
Glucose flux in lean and *ob/ob* mice primary hepatocytes. **a,** Schematic illustration of isotope tracing experiments with [U-^13^C]Glc in mouse primary hepatocytes. Mouse primary hepatocytes were isolated from lean, wild-type and *ob/ob* mice, incubated with minimal medium or complete medium (glucose-free DMEM) for 45 minutes, then replaced with corresponding medium plus 10 mM [U-^13^C]Glc tracer, and terminated either after 2, 5, 10, and 30 minutes (in minimal medium) or after 10 and 60 minutes (in DMEM). Glucose, Glc. **b-j,** Accumulation kinetics of uniform,^13^C isotope-labeled glucose and metabolites in lean and *ob/ob* mice primary hepatocytes during the course of [U-^13^C]Glc tracing. Glyceraldehyde-3-phosphate,GA3P; dihydroxyacetone phosphate, DHAP; phosphoenolpyruvate, PEP. **k,** Heatmap of glucose metabolite profile from the liver tissues of lean and *ob/ob* mice. Color scale denotes row scaling z-scores. Glucose 6-phosphate/fructose 6-phosphate, G6P/F6P; fructose 1,6-diphosphate, FBP; 1,3-bisphosphoglycerate, 1,3-BPG; 3-phosphoglycerate/2-phosphoglycerate, 3PG/2PG; pyruvate, Pyr; lactate, Lac. **l,** Fractional abundance of ^13^C-labeled glycolysis intermediates compared to the total (labeled and unlabeled) of corresponding metabolites in lean and *ob/ob* primary hepatocytes at steady state (30 minutes). **m,** Fractional abundance of all ^13^C-labeled TCA intermediates at steady state. Abbreviations: Acetyl-CoA, Ac-CoA; oxaloacetate, OAA; citrate, Cit; aconitate, Aco; isocitrate, IsoCit; α-ketoglutarate, α-KG; succinate, Succ; fumarate, Fum; malate, Mal. (N) Accumulation kinetics of ^13^C-labeled succinate in lean and *ob/ob* mice primary hepatocytes. **o,** Measurement of glucose-induced change in mitochondria oxygen consumption rate (OCR) in lean and *ob/ob* primary hepatocytes (n=8 and 6 respectively). **p,** Illustration of obesity-induced hepatic metabolic remodeling inferred from [U-^13^C]Glc tracing studies. Data are presented as means ± SEM, n=3; text labeled 0.05 < p < 0.1, *p < 0.05, **p < 0.01, ***p < 0.001, two-tailed unpaired Student’s *t*-test (*ob/ob vs.* lean). N.D., not detected. See also Supplementary Figure 1 and 2.

Primary hepatocytes were first isolated from the wild-type, lean and *ob/ob* mice and fasted for 45 minutes to deplete intracellular glucose and glycogen (Supplementary Fig. 1a). A 10mM, ^13^C-labeled glucose was then added into a fresh tracing medium, and samples were collected at a series of time points after medium replacement, analyzed by liquid chromatography-mass spectrometry (LC-MS). The uptake of glucose was rapid and plateaued after two minutes (Fig. 1b). At the 10-minute time point, the upper part of glycolysis also reached a plateau (Fig. 1c, d; Supplementary Fig. 1b), while the lower part continued to increase toward the 30-minute mark (Fig. 1e-g; Supplementary Fig. 1c). Lactate and glycerol-3-phosphate are two major carbon sinks for glycolysis, and both of them were accumulated 2-3 times faster in the obese mice primary hepatocytes than in the lean (Fig. 1f-h). Intriguingly, the PPP metabolite ribose-5-phosphate (R5P) was accumulated faster in the lean than in the obese primary hepatocyte (Fig. 1i), possibly due to hyperactivation of the downstream non-oxidative PPP (sedoheptulose-7-phosphate, S7P, Fig. 1j) and purine synthesis pathway (inosine-5-monophosphate, IMP, Supplementary Fig. 1d). Fluxing the primary hepatocytes in a complete medium supplemented with amino acids reduced the fractional labeling of many of the metabolites, but the overall difference between lean and obese primary hepatocytes remained (compare Supplementary Fig. 1b-h in minimal medium to Supplementary Fig. 1i-o in complete medium), suggesting that the nutritional status did not perturb the endogenous metabolic program in a significant way. The authenticity of flux analysis was further corroborated by the observation that glycolytic metabolites from the obese liver tissues are uniformly higher than from the lean tissues (Fig. 1k).

Despite the near complete labeling of pyruvate by glucose (Fig. 1l), the labeling efficiency of acetyl-CoA and TCA cycle intermediates was very low (Fig. 1m, n; Supplementary Fig. 1q-s), suggesting the pyruvate dehydrogenase (PDH) activity being rate-limiting. Nonetheless, the expected effect of hyperinsulinemia on PDH was preserved in the obese primary hepatocytes, as glucose labeling of the majority of TCA cycle intermediates trended upwards (Fig. 1m, n). By measuring the oxygen consumption rate (OCR), we were able to confirm that mitochondrial glucose oxidation was indeed increased in the obese primary hepatocyte (Fig. 1o).

Therefore, the development of obesity seems to increase all aspect of glucose catabolism (Fig. 1p). The hyperactivation of glucose catabolism exerts spill-over effects by enhancing the *de novo* synthesis of non-essential amino acids (Supplementary Fig. 1h), and it reduces amino acid catabolism toward the gluconeogenic and glyceroneogenic pathways (Fig. 1p).

### Glutamine flux concentrates around the TCA cycle

To directly assess the fate and flux of amino acid metabolism, we traced the uniformly labeled glutamine ([U-^13^C, ^15^N]Gln) in both lean and obese primary hepatocytes at 2.5mM concentrations (Fig. 2a).

**Figure 2.**
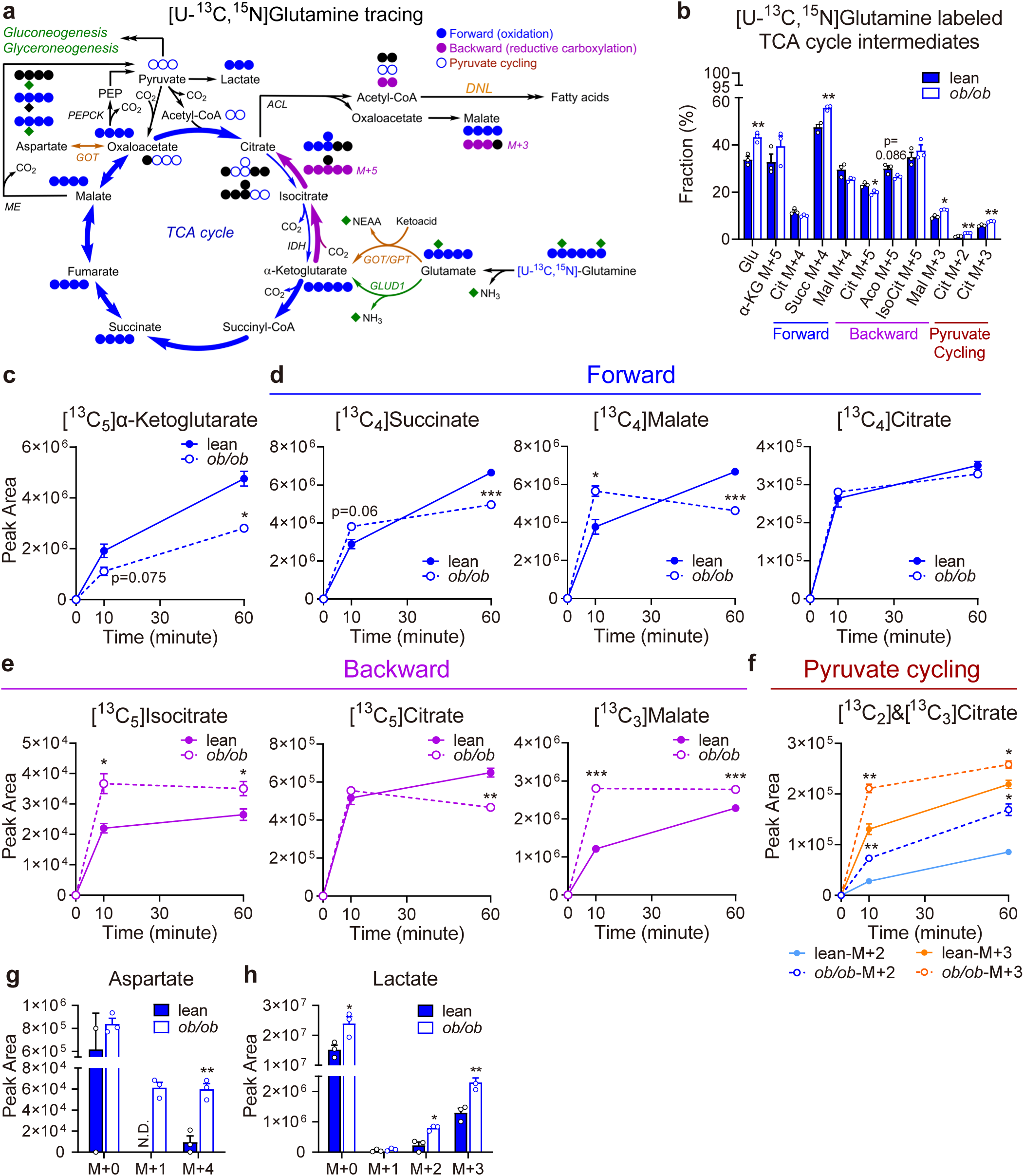
Glutamine flux in lean and *ob/ob* mice primary hepatocytes. **a,** Schematic illustration of [U-^13^C, ^15^N]Gln tracing. Primary hepatocytes were first primed in M199 or minimal medium for 45 minutes, then replaced with corresponding medium containing 2.5 mM tracer, and terminated either after 10 and 60 minutes (in M199) or after 5, 10, and 30 minutes (in minimal medium). Gln, glutamine; GOT: glutamate-oxaloacetate transaminase; GPT: glutamate-pyruvate transaminase; GLUD1: glutamate dehydrogenase 1; IDH: isocitrate dehydrogenase; ACL: ATP citrate lyase; ME: malic enzyme; PEPCK: phosphoenolpyruvate carboxykinase. **b,** Fractional abundance of all ^13^C isotope-labeled TCA cycle intermediates compared to the total (labeled and unlabeled) of corresponding metab- olite in the primary hepatocytes 10 minutes after the addition of by [U-^13^C, ^15^N]Gln. Glutamate, Glu; oxaloacetate, OAA; citrate, Cit; aconitate, Aco; isocitrate, IsoCit; α-ketoglutarate, α-KG; succinate, Succ; malate, Mal. **c-f,** Accumulation kinetics of ^13^C isotope-labeled TCA cycle intermediates during the course of [U-^13^C, ^15^N]Gln tracing. **g,** Abundance of ^13^C (M+4) or ^15^N (M+1) isotopomers of aspartate accumulated in primary hepatocytes 10 minutes after [U-^13^C, ^15^N]Gln addition. **h,** Abundance of isotopomers of ^13^C-labeled (M+1, 2, 3) and unlabeled (M+0) lactate accumulated at 10 minutes after [U-^13^C, ^15^N]Gln addition. Data are presented as means ± SEM, n=3; text labeled 0.05 < p < 0.1, *p < 0.05, **p < 0.01, ***p < 0.001, two-tailed unpaired Student’s *t*-test (*ob/ob vs.* lean). N.D., not detected. See also Supplementary Figure 3.. **b,** Fractional abundance of all ^13^C isotope-labeled TCA cycle intermediates compared to the total (labeled and unlabeled) of corresponding metabolite in the primary hepatocytes 10 minutes after the addition of by [U-^13^C, ^15^N]Gln. Glutamate, Glu; oxaloacetate, OAA; citrate, Cit; aconitate, Aco; isocitrate, IsoCit; α-ketoglutarate, α-KG; succinate, Succ; malate, Mal. **c-f,** Accumulation kinetics of ^13^C isotope-labeled TCA cycle intermediates during the course of [U-^13^C, ^15^N]Gln tracing. **g,** Abundance of ^13^C (M+4) or ^15^N (M+1) isotopomers of aspartate accumulated in the primary hepatocytes 10 minutes after the addition of [U-^13^C, ^15^N]Gln. **h,** Abundance of all isotopomers of ^13^C-labeled (M+1, 2, 3) and unlabeled (M+0) lactate accumulated in the primary hepatocytes 10 minutes after the addition of [U-^13^C, ^15^N]Gln. **i,** Illustration of obesity-induced hepatic metabolic remodeling inferred from [U-^13^C, ^15^N]Gln tracing studies. Line thickness represents the relative flux rate of different pathways. Data are presented as means ± SEM, n=3; text labeled 0.05 < p < 0.1, *p < 0.05, **p < 0.01, ***p < 0.001, two-tailed unpaired Student’s *t*-test (*ob/ob vs.* lean). N.D., not detected. See also Supplementary Figure 3..

Glutamine is first metabolized to glutamate through glutaminolysis (Fig. 2b, Supplementary Fig. 3a-c) and then converted to α-ketoglutarate (α-KG) through either transamination or oxidative deamination pathways (Fig. 2b, c); α-ketoglutarate may then enter TCA cycle for oxidation in the mitochondria (forward, Fig. 2d), or go through reductive carboxylation (backward, Fig. 2e) for the synthesis of citrate toward DNL in the cytosol. The carbon backbones of glutamine entering into TCA cycle may also exit at the level of oxaloacetate (OAA), which may re-enter the TCA cycle through pyruvate cycling (Fig. 2f), be converted into aspartate (Asp) through transamination reactions (Fig. 2g), or participate in glyceroneogenesis/gluconeogenesis pathways (Fig. 2h).

The supplement of 2.5mM glutamine ([U-^13^C, ^15^N]Gln) in the tracing medium led to a rapid appearance and dominance of labeled glutamine and glutamate in the intracellular pool (Supplementary Fig. 3b, c). Although the 2.5mM glutamine concentration used herein is higher than the physiological level of glutamine in circulation, it only caused a transient spike in the intracellular glutamine pool, and no elevation in glutamate concentration was observed, suggesting the presence of an inherent mechanism in maintaining cellular glutamine levels.

The labeled glutamine carbon then rapidly entered the TCA cycle and plateaued at ∼10 minutes (Fig. 2d, e). The continued accumulation of α-KG after the 10 minute time point (Fig. 2c) suggests the combined activity of glutamate transamination and oxidative deamination pathways exceeded α-KG consumption rates. At steady state, the fraction of TCA metabolites labeled by glutamine-derived carbon reached ∼40%, 5-10 times higher than that of glucose (Fig. 2b). The fate of glutamine metabolism was similar when the primary hepatocytes were cultured under either complete (Fig. 2c-h, Supplementary Fig. 3a) or minimal media conditions (Supplementary Fig. 3b-i). But the TCA intermediate accumulation was slower in the minimal medium, suggesting the presence of glucose suppressed cataplerosis of glutamine catabolite from TCA.

The development of obesity hastened the kinetics of glutamine entrance into the TCA cycle (Fig. 2d-f), although the accumulation of α-KG was slower in the obese primary hepatocytes (Fig. 2c), again confirming α-KG production was not rate-limiting in glutamine catabolism. The intermediates generated from reductive carboxylation direction (backward) was accumulated more significantly than forward TCA metabolites (Fig. 2e, Supplementary Fig. 3f). The effect of obesity on glutamine catabolism pathways entering and exiting the TCA cycle, however, differed significantly. As shown in Fig. 2g and Supplementary Fig. 3j, the synthesis of aspartate, either from glutamate derived carbon backbone (M+4) or donated amine group (M+1), increased sharply in the obese primary hepatocytes, suggesting an upregulated transamination to facilitate glutamine catabolism. Similarly, the returning of OAA carbon toward aerobic glycolysis (lactate, Fig. 2h) or TCA cycle (pyruvate cycling, Fig. 2g, Supplementary Fig. 3g) through the cataplerosis pathway was increased. In contrast, the synthesis of glycerol-3P from glutamine through the glyceroneogenic route decreased in the obese primary hepatocytes (Supplementary Fig. 3i). The reduction in glutamine glyceroneogenesis coincided with induction of glycolysis in the obese primary hepatocytes, suggesting a partial replacement of glutamine footprint by glucose in the obese state. Therefore, unlike glucose, the carbon flux of glutamine is concentrated in and around the TCA cycle, and the metabolic profile of glutamine carbon is further compressed in the obese primary hepatocytes through a coordinated metabolic reprogramming that results in an overall reduction in glyceroneogenesis.

**Figure 3.**
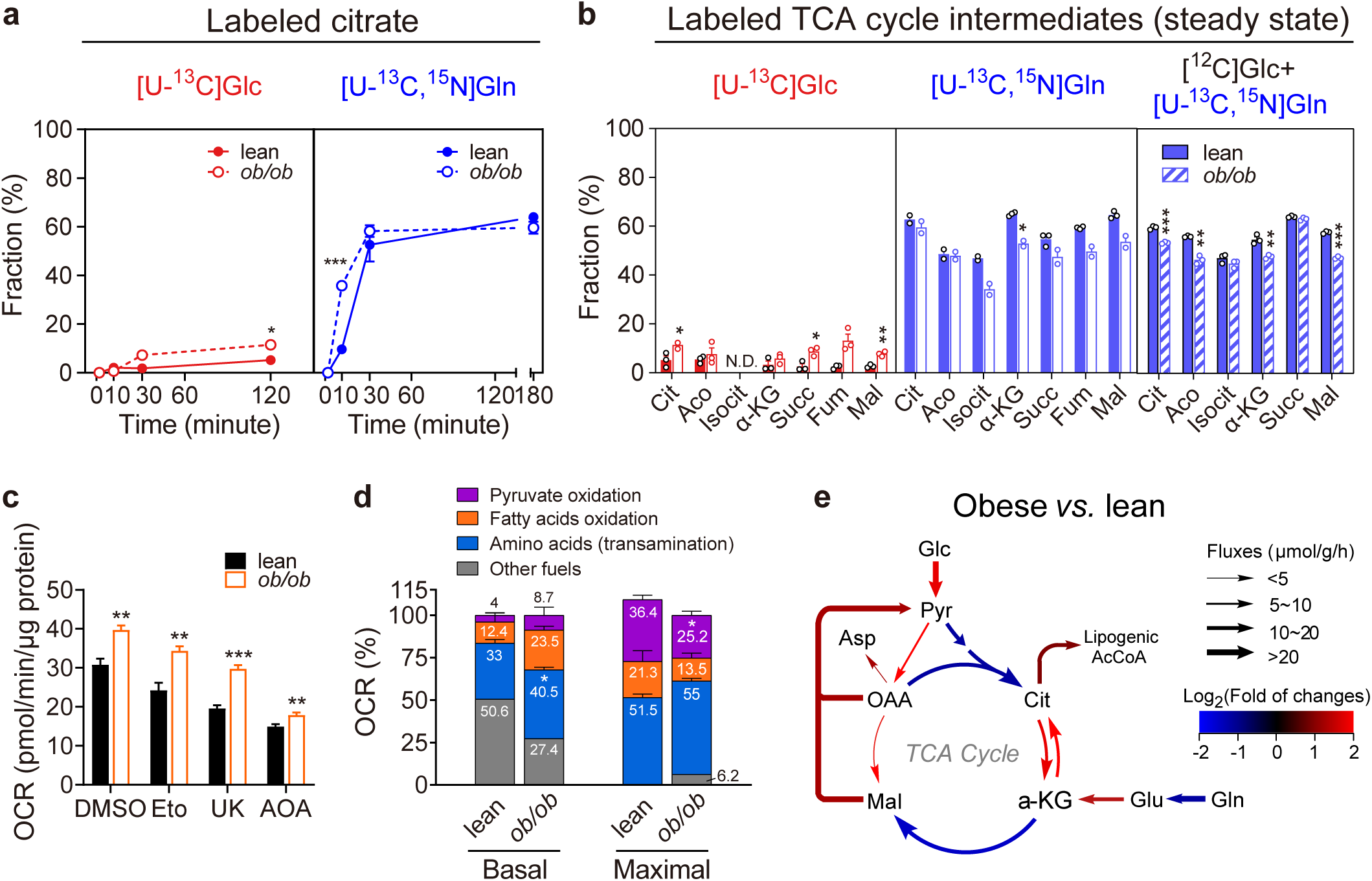
Substrate level contribution of glucose and glutamine toward TCA cycle. **a,** Kinetics of ^13^C isotope-labeled citrate accumulation in lean and obese primary hepatocytes traced with either 10 mM [U-^13^C]Glc or 2.5 mM [U-^13^C, ^15^N]Gln. **b,** Fractional abundance of all ^13^C isotope-labeled TCA intermediates compared to the total (labeled and unlabeled) of corresponding metabolites at steady state. Lean and *ob/ob* primary hepatocytes were traced with either 10 mM [U-^13^C]Glc or 2.5 mM [U-^13^C, ^15^N]Gln in minimal medium, or 2.5 mM [U-^13^C, ^15^N]Gln in M199 ([^12^C]Glc + [U-^13^C, ^15^N]Gln). **c,** Maximal mitochondrial oxygen consumption rate (OCR) of lean and *ob/ob* primary hepatocytes treated with DMSO (0.1%), etomoxir (Eto, 40 μM), UK5099 (UK, 40 μM), or aminooxyacetate (AOA, 40 mM). **d,** Substrate-specific contribution toward mitochondria respiration calculated from (**c**). **e,** Graphical summary of TCA cycle fluxes estimated by metabolic flux analysis of the data in Figure 1-3. Flux rate are expressed as μmol/g cells per hour. Line colors represent the flux changes between lean and obese mice primary hepatocytes. Abbreviations: glucose, Glc; pyruvate, Pyr; glutamine, Gln; glutamate, Glu; oxaloacetate, OAA; citrate, Cit; acetyl-CoA, Ac-CoA; α-ketoglutarate, α-KG; malate, Mal; aspartate, Asp. See details in Supplementary Figure 4g and Table 1. Data are presented as means ± SEM, outliers beyond 1.5×SD are removed; text labeled 0.05 < p < 0.1, *p < 0.05, **p < 0.01, ***p < 0.001, two-tailed unpaired Student’s *t*-test (*ob/ob vs.* lean). N.D., not detected. See also Supplementary Figure 4.

### Glutamine, not glucose, is the principle substrate for oxidative respiration in hepatocytes

The flux analyses above revealed distinct rate-limiting steps of glucose and glutamine metabolism toward the TCA cycle: PDH for glucose and α-ketoglutarate consumption for glutamine. As a result, albeit the kinetics of TCA intermediates labeling by glucose and glutamine did not differ significantly (Fig. 3a; Supplementary Fig. 1q-s, Supplementary Fig. 3d, e; Supplementary Fig. 4a), their steady-state labeling efficiency differed for about tenfold (less than 5% by glucose *vs.* ∼50% by glutamine, Fig. 3b; Supplementary Fig. 4b).

Oxidative phosphorylation in the liver primarily depends on complex II of the electron transport chain^24^, and glutamine catabolism is proximally positioned toward complex II in four enzymatic steps. However, amino acid was not considered as the main substrate for energy production. To independently measure substrate contribution toward the TCA cycle, we compared mitochondria respiration in the presence or absence of inhibitors for transaminases (aminooxyacetate, AOA), pyruvate transporters (UK5099, UK), and fatty acid transporters (Etomoxir, Eto) (Fig. 3c, Supplementary Fig. 4c-f). As quantified in Fig. 3d, the amino acid (33%) is the single most dominant substrate oxidized by mitochondria, followed by fatty acids (12.4%) and pyruvate (4%, used as a surrogate for glucose). The contribution of pyruvate toward mitochondria respiration was doubled to 8.7% in the obese primary hepatocytes (Fig. 3d), similar to the glucose labeling efficiency of TCA intermediates under the same condition (Fig. 3b). Only after the addition of the mitochondria respiration uncoupling reagent, pyruvate may enter the TCA cycle in a significant amount (36.4%, Fig. 3d, maximal), possibly due to the removal of product (NADH) inhibition at the level of PDH.

We further applied OpenMebius flux modeling^25^ to understand the interaction between glucose and glutamine metabolism and how obesity modifies their interaction. Modeling analysis confirmed glutamine as a larger contributor to TCA cycle than glucose. The calculated rate of glycolysis (glucose to pyruvate) in lean mice primary hepatocytes was only 1/2 to 1/3 of the rate of glutamine catabolism to α-ketoglutarate (Fig. 3e; Supplementary Fig. 4g; Supplementary Table 1). By comparing flux rates in glucose and glutamine tracing in the minimal medium, we further identified a two-way interaction between glucose and glutamine metabolism. On the one hand, glucose stimulated glutamine utilization through glutaminolysis, oxidation, and reductive carboxylation (Supplementary Table 1). On the other hand, glutamine suppressed glycolysis and pyruvate oxidation but increased lactate production (Supplementary Table 1). The development of obesity had distinct effects on glucose and glutamine metabolism: it tripled the rate of glycolysis (glucose-pyruvate increased from 6.27 to 19.88μmol/g/min), and more than quadrupled (from 6.89 to 29.81μmol/g/min) the reductive carboxylation of glutamate, but the rate of glutamate oxidation was significantly reduced (from 17.85 to 6.76 μmol/g/min; Supplementary Fig. 4g; Supplementary Table 1). Together, these results suggest glucose stimulates glutamine utilization while glutamine complements (through gluconeogenesis) and competes (by preventing pyruvate entrance into TCA) with glucose metabolism. The development of obesity increases the overall flux by elevating gluconeogenesis, lipogenesis and lactate production.

### Glutamine is a prominent carbon supplier for hepatic DNL

Citrate synthesized in the TCA cycle may be used for substrate oxidation or lipogenesis. The revelation that amino acid but not glucose is the main substrate of oxidative phosphorylation led us to review another long-held view that carbohydrates drive lipogenesis. We applied similar, side-by-side comparative flux analysis of glucose and glutamine toward the synthesis of triglyceride fatty acid in primary hepatocytes isolated from overnight-fed, lean and obese (*ob/ob*) mice (Fig. 4a). We chose high-glucose (15mM), high-glutamine (3.2mM), and high-insulin conditions with a physiological level (hepatic portal vein) of acetate (0.61mM) to sustain lipogenesis *ex vivo*^26, 27^. The duration of labeling was extended to 6 hours to increase labeling efficiency, without compromising the expression of key enzymes (Supplementary Fig. 5a). The overall fatty acid pool was also reflective of its lipogenic signature in that the primary DNL products of palmitate (C16:0) and oleate (C18:1) were highly elevated in the obese primary hepatocyte compared to that of lean, whereas the essential fatty acids including linoleic (C18:2) and arachidonic acid (C20:4) did not differ between lean and obese primary hepatocytes (Supplementary Fig. 5b).

**Figure 4.**
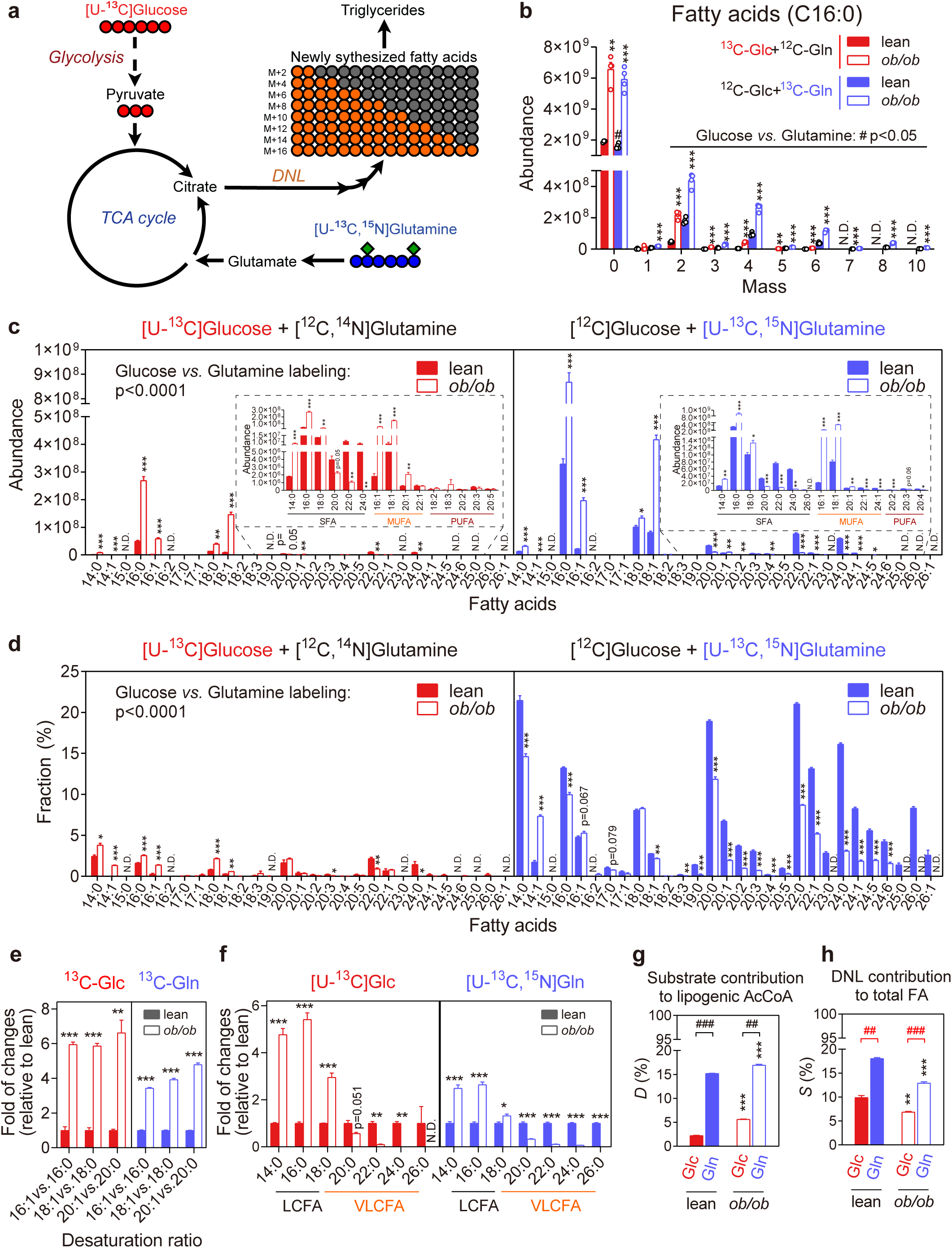
Carbon backbone flux from glucose and glutamine toward fatty acids. **a,** Schematic illustration of glucose and glutamine carbon backbone flux toward fatty acid (FA) synthesis. Lean and *ob/ob* primary hepatocytes were primed with M199 and 10 nM insulin for 45 minutes. Then the culture medium was replaced with M199 and 10 nM insulin supplemented with either 10 mM [U-^13^C]Glc plus 2.5 mM unlabeled Gln or 2.5 mM [U-^13^C, ^15^N]Gln plus 10mM unlabeled Glc for 6 hours. M199 contained 0.61mM acetate and other essential nutrients for hepatocytes culture. **b,** Abundance of triglyceride palmitate (C16:0 FA) isotopomers in lean and *ob/ob* primary hepatocytes traced with either 10 mM [U-^13^C]Glc or 2.5 mM [U-^13^C, ^15^N]Gln as described in (**a**). **c**,**d,** Normalized (**c**) and fractional (**d**) abundance of ^13^C-labeled triglyceride fatty acid isotopomers in primary hepatocytes traced with either 10 mM [U-^13^C]Glc or 2.5 mM [U-^13^C, ^15^N]Gln. The abundance of isotopomers with odd numbers of ^13^C was very low and excluded from all calculations. SFA, saturated fatty acids; MUFA, monounsaturated fatty acids; PUFA, polyunsaturated fatty acids. **e,** Relative abundance of ^13^C-labeled even chain MUFA compared to corresponding SFA in lean and *ob/ob* primary hepatocytes. **f,** Relative abundance of all ^13^C-labeled FA isotopomers compared to the total amount of corresponding FA in lean mice primary hepatocytes. LCFA, long chain fatty acids of C16 and C18; VLCFA, very long chain fatty acids of C20-26. **h**, **i**, Fractional contribution of ^13^C-labeled acetyl-CoA (AcCoA) (*D*%, **h**) toward the total lipogenic AcCoA pool, and the contribution of *de novo* synthesized fatty acid (*S*%, **i**) during the period of tracing studies toward total fatty acid pool. *D*% and *S*% are calculated through the isotopomer aggregation method (see materials and methods) using isotopomer fractional abundance and ^13^C mole percent enrichment (MPE) of palmitate (C16:0 FA) as input (see Supplementary Figure 5g). Red line and # text labeling denote the estimation bias of *S*%. Please refer to Supplementary Figure 7j for comparison. Data are presented as means ± SEM, n=4 except that one sample was discarded in the ^13^C Glc-labeled lean group; text labeled 0.05 < p < 0.1, *p < 0.05, **p < 0.01, ***p < 0.001 (*ob/ob vs.* lean) and #p<0.05 or text labeled (Glc *vs.* Gln) by two-tailed unpaired Student’s *t*-test; text labeling on the top left of panel **c-d** are significant based on two-way ANOVA (Glc *vs.* Gln). N.D., not detected. See also Supplementary Figure 5, 6, 7.

Consistent with their distinct efficiency in citrate labeling, glutamine but not glucose-derived ^13^C was readily incorporated into palmitate in the lean mice primary hepatocytes (Fig. 4b; Supplementary Fig. 5c). Notably, isotopic enrichments of top ranked FA from glutamine (^13^C_5_, ^15^N_2_) ranged from 5%-22%, overwhelmingly greater than the 0.2-2.6% enrichment of the glucose-derived ^13^C carbons (Fig. 4d). Glucose incorporation increased by fivefold in the obese primary hepatocytes (Fig. 4b, c), more than the overall increase of glycolysis and citrate labeling under the same conditions. As a result, the fractional contribution of glucose to palmitate synthesis was almost doubled in the obese primary hepatocytes (Fig. 4d, Supplementary Fig. 5c) despite an almost three-fold increase in the total triglyceride-palmitate levels (Fig. 4b). The obesity-induced increase of glutamine flux into palmitate was not as substantial (Fig. 4b-d), and its fractional contribution was reduced (Fig. 4d; Supplementary Fig. 5c). Still, the overall contribution of glutamine toward palmitate synthesis remained about three times higher than that of glucose (Fig. 4d).

Oleate (C18:1), palmitoleate (C16:1), stearate (C18:0) are immediate elongation and desaturation products of palmitate, and all of them were adequately labeled (Fig. 4c, d; Supplementary Fig. 5d-f). However, the ratio of newly synthesized unsaturated *vs.* saturated fatty acids (C16:1/C16:0 and C18:1/C18:0) was higher in the obese primary hepatocytes (Fig. 4e), suggesting increased desaturation in the obese than the lean. In contrast, the ability of the obese primary hepatocytes to elongated fatty acids was significantly compromised. The lean mice primary hepatocytes synthesized more C20-24 than their obese counterparts (Fig. 4f) despite their much lower DNL activity. Consequently, the overall composition of very long chain fatty acids (VLCFA) was reduced in the obese primary hepatocytes (Supplementary Fig. 5g). A mixture of amino acids other than glutamine was also able to readily label palmitate in the primary hepatocytes, suggesting amino acids as a whole are important DNL substrates (Supplementary Fig. 5h), and a similar deficiency in fatty acid elongation was observed in obese primary hepatocytes in mixed amino acid tracing experiments (Supplementary Fig. 5i).

The preferred incorporation of glutamine over glucose carbon into DNL products was not affected by dietary conditions, as primary hepatocytes isolated from high-fat diet (HFD)-fed mice exhibited a similar preference (Supplementary Fig. 6). Nor was it driven by high glutamine concentration in the tracing medium either. As shown in Supplementary Fig. 7a-c, regardless of culture medium glutamine concentration, the intracellular glutamine concentration of primary hepatocytes remained significantly lower than that of liver tissues, whereas the glutamate pool was modestly higher. Additionally, an increase of glutamine concentration in the culture medium from 0.5mM to 2.5mM did not suppress glucose entry into the DNL pathway at the level of pyruvate dehydrogenase nor induce lipogenic enzyme expression (Supplementary Fig. 7d, e). This is further supported by the observation that changes in tracing glutamine concentration affected the labeling efficiency of intracellular glutamine (Supplementary Fig. 7b) and fatty acid pool (Supplementary Fig. 7f), but not the overall contribution of glutamine toward DNL (Supplementary Fig. 7g).

Current computational methods do not work well for short-term, low-efficiency labeling experimental systems^28–30^, and primary hepatocytes studied herein are not amenable to long-term studies due to their propensity for de-differentiation. A major reason is fitting background level mass isotopomer distribution (MID) led to arbitrarily high *D* values and compressed *S* values^28^. For example, the percentile of newly synthesized palmitate (*S*%) calculated by the Fatty Acid Source Analysis (FASA)^31^ in the ^13^C-glucose trace experiment was a mere 0.04%, far lower than the 1.64% of newly synthesized palmitate containing ^13^C- glucose-derived C2 units (mass isotopomer enrichment, MIE, Supplementary Fig. 7h), which should be a fraction of the former.

To avoid over-fitting of the background level mass isotopomer distribution signal, we opted to aggregate isotopomer signals to estimate the fractional contribution of the labeled substrate as well as the rate of DNL (Fig. 4g and h; see “Mass Isotopomer Enrichment (MIE)-based Estimate of DNL” in the Methods section). Through this method, we estimated that the fractional contribution (*D*%) of ^13^C-labeled glutamine toward palmitate was between 15∼17% in the lean mice primary hepatocytes as well as the HFD and *ob/ob* mice, all of which were much higher than those of ^13^C-labeled glucose (Fig. 4g; Supplementary Fig. 7i). Increasing ^13^C-glucose concentration from 10mM to 20mM boosted the fractional contribution of glucose to ∼9% (Supplementary Fig. 7j), suggesting a non-linear contribution under hyperglycemia conditions.

The amount of palmitate synthesized (*S*%) during the 6h tracing period was estimated to be ∼10% of the total palmitate pool, and it was slightly lower in the obese primary hepatocytes than in the lean (Fig. 4h), reflective of their much larger intracellular triglyceride pool. Importantly, the difference between the estimated *S*% under glucose and glutamine tracing conditions was much smaller compared to that of the FASA method (compare Fig. 4h and Supplementary Fig. 7j), another indication that the MIE method overperforms MID for low-efficiency labeling studies.

### Quantitative contribution of glucose and amino acid carbon toward lipogenesis *in vivo*

Metabolism does not occur in isolation, and dietary nutrients may contribute to hepatic metabolism indirectly after initial transformation in other tissues^32, 33^. Therefore, we conducted dietary glucose and amino acid tracing through drink to examine the physiological absorption and metabolism of nutrients *in vivo* (Fig. 5a-j; Supplementary Fig. 8a-i).

**Figure 5.**
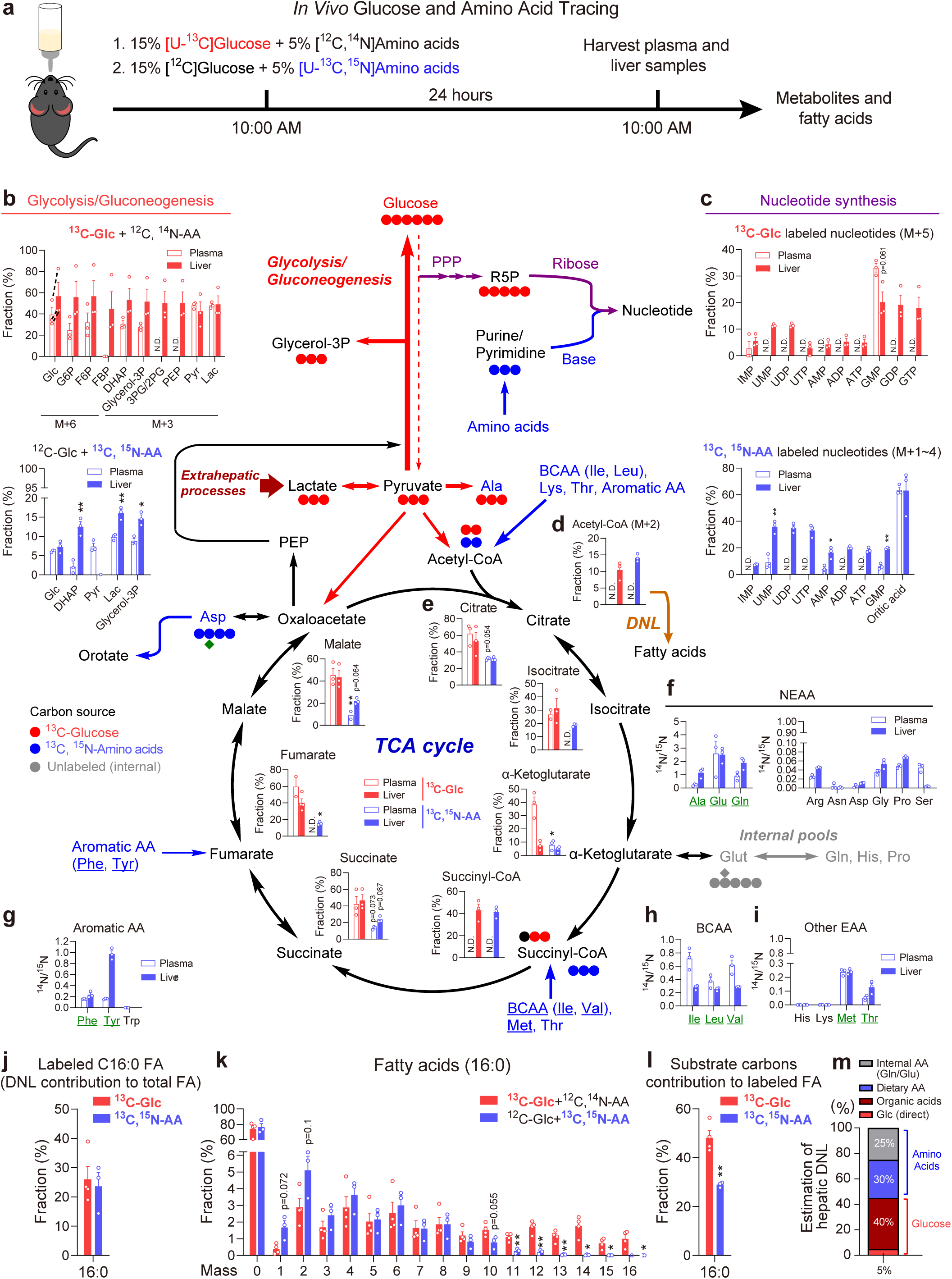
*In vivo* metabolic flux of glucose and amino acids. **a,** Schematic illustration of experimental design for *in vivo* glucose and amino acid isotope tracing. 12-week-old mice were fed with drinking water containing either 15% [U-^13^C]glucose plus 5% unlabeled amino acid mixture or 15% unlabeled glucose plus 5% [U-^13^C,^15^N]amino acid mixture for 24 hours. **b, c,** Fractional abundance of ^13^C labeled glycolysis/gluconeogenesis (**b**) and nucleotide synthesis (**c**) intermediates in plasma (open bar) and liver (solid bar) traced with either [U-^13^C]glucose (upper) or [U-^13^C, ^15^N]amino acid mixture (lower).. **d, e,** Fractional abundance of ^13^C labeled liver acetyl-CoA (**d**) and TCA cycle intermediates (**e**) traced with either [U-^13^C]glucose or [U-^13^C, ^15^N]amino acid mixture. **f, g, h, i,** The ^13^C^14^N/^13^C^15^N ratio of amino acids indicating catabolic rate (catabolism *vs.* reamination): NEAA (except tyrosine, **f**), aromatic AA (**g**), branched chain amino acids (BCAA, **h**), and other EAA (**i**). AA with high ^13^C^14^N/^13^C^15^N ratio (>0.1) are labeled with underlined green texts. **j, k,** Total (**j**) and separative (**k**) fractional abundance of ^13^C labeled liver triglyceride palmitate (fatty acid 16:0) isotopomers. **l,** Fractional abundance of the labeled carbons in the newly synthesized palmitate (**j**). **m,** Schematic illustration for the estimation of hepatic DNL substrates composition. Data are presented as means ± SEM; text labeled 0.05 < p < 0.1, *p < 0.05, **p < 0.01, ***p < 0.001 (plasma *vs.* liver or Glc *vs.* AA). N.D., not detected. See also Supplementary Figure 8.

We first examined glucose absorption and metabolism in mice fed with 15% uniformly labeled, ^13^C-glucose ([U-^13^C]Glc) supplemented with 5% unlabeled amino acid mixture (Fig. 5a; Supplementary Fig. 8a). Mass spectrometry analysis showed that labeled glucose were efficiently transformed into organic acids in the plasma and the liver (Fig. 5b, e; Supplementary Fig. 8b, c), consistent with previous observations made by the Rabinowitz group that carbohydrates were first transformed into organic acids and released into circulation before being utilized for energy production^32, 33^. Besides supplying the TCA cycle (Fig. 5e), organic acids were fed into hepatic gluconeogenesis as indicated by the enrichment of ^13^C-labeled gluconeogenic intermediates and glucose isotopomers in liver over their plasma counterparts (Fig. 5b). Dietary glucose further contributed to the hepatic synthesis of the ribose group through the pentose phosphate pathways as evidenced by the m+5 and m+10 labeling of nucleotides (m+5, Fig. 5c upper panel). However, most amino acids, except alanine (Supplementary Fig. 8f), were not labeled by glucose-derived ^13^C, suggesting a weak cataplerosis from the TCA cycle. Together, these data suggest dietary glucose utilization in the liver is facilitated by its initial transformation into organic acids in extra-hepatic tissues, and hepatocytes themselves do not catabolize glucose for energy production or non-essential amino acid synthesis.

We then examined amino acid metabolism in mice fed with uniformly labeled amino acid mixture ([U-^13^C, ^15^N]AA) in the presence of unlabeled glucose (Supplementary Fig. 8a-e). Overall, amino acid as a mixture were metabolized very differently *in vivo*. For example, glutamine, glutamate, and aspartate were effectively cleared by the intestine, and the labeled forms were almost absent from the plasma (Supplementary Fig. 8e, f). In contrast, the majority of essential amino acids were absorbed into the plasma efficiently, and lysine was uniquely enriched in the plasma (Supplementary Fig. 8e), possibly due to its superfluous composition in the mixture (Supplementary Fig. 8a). The carbon backbones of those cleared amino acids were transformed into ^13^C-labeled organic acids, including lactate, pyruvate, citrate, succinate, and orotate (Fig. 5b-c lower panel, e), which may serve as precursors for energy production, gluconeogenesis, and nucleotide biosynthesis (Fig. 5c lower panel). Therefore, the initial transformation of dietary nutrients into organic acids is not a carbohydrate-specific phenomenon. However, labeled amino acid carbon did not enter the gluconeogenic pathways effectively (Fig. 5b lower panel), reaffirming the territorial dominance of glucose carbon in this domain. Additionally, the absence of labeled α-ketoglutarate in the plasma suggests the carbon backbones of glutamine and glutamate might be utilized by the intestinal microbiota (Fig. 5e), and the body glutamine/glutamate pool is most likely synthesized from other organic acids.

The difference in amino acid catabolism may also be inferred from the relative abundance of [U-^13^C, ^14^N] AA over their respective [U-^13^C, ^15^N] isotopomers. As shown in Fig. 5g-j, the ^13^C^14^N/^13^C^15^N ratio is closely correlated with the labeling efficiency of the TCA cycle metabolites generated from their respective amino acids (underlined). Aspartate is the only exception in that it has no detectable levels of ^13^C^14^N-Asp (Fig. 5j right panel), because the donation of its amine group for adenine synthesis from IMP (Fig. 5c) is accompanied by the release of its carbon backbone as formate, preventing re-amination. Importantly, the labeling efficiency of acetyl-CoA were higher in the amino acid-traced samples than in the glucose-traced samples (Fig. 5d), despite glucose being three times higher in abundance than the whole amino acid mixture, and dietary non-essential amino acids were effectively cleared by the intestine.

Lastly, we examined how dietary glucose and amino acids labeled the hepatic fatty acid pool under above tracing conditions (Fig. 5a). As shown in Fig. 5k, the percentage of fatty acids with ^13^C incorporation varied significantly both among different fatty acid species and within each experimental group (Fig. 5j; Supplementary Fig. 8g), suggesting individual variations among hepatic DNL activity. However, the fatty acid isotopomer relative abundance distribution curve was remarkably similar within each group (Fig. 5k; Supplementary Fig. 8h). The abundance of glucose-labeled palmitate and stearate isotopomer species were relatively equivalent, whereas the amino acid labeled isotopomer distribution curves were strongly skewed toward low m+ isotopomers (Fig. 5k), indicating higher carbon contribution from glucose than from amino acids. However, the complex, zig-zagging isotopomer distribution pattern between odd and even m+ isotopomers suggests a non-homogenous polymer synthesis process, and it prevents the exact determination of the fractional contribution of glucose and amino acid derived carbon toward DNL based on mass isotopomer analysis^30^. Instead, we aggregated the percentage of ^13^C among the labeled fatty acids as an approximation of fractional substrate contribution. As shown in Fig. 5l, dietary glucose contributed ∼45% of carbon toward newly synthesized palmitate, while amino acids contributed ∼30%. Given the amount of amino acids was only 1/3 of glucose in the drinking water, the lipogenic potential of dietary amino acids is ∼2-fold greater than that of glucose. The fractional contribution of dietary glucose and amino acid carbon toward the synthesis of other fatty acids in the liver was similar to those of palmitate (Supplementary Fig. 8i), and the overall contribution of glucose and amino acids toward hepatic DNL was around 45% *vs.* 55% when we take the endogenous amino acid pool into account (Fig. 5m).

### Stimulation of glycogen storage reduces glycerol-3P synthesis and hepatic TAG accumulation

The *ex vivo* and *in vivo* flux analyses suggest glucose may contribute to triglyceride synthesis through the synthesis of glycerol-3P and fatty acids. We reasoned that an increase in glycogen synthesis may serve as a sink to compete glucose and its organic acid derivatives away from triglyceride synthesis pathways in obesity.

An initial support for this hypothesis was from the observation that the obese mice exhibited prolonged hyperglycemia after food ingestion and a severed ability to synthesize hepatic glycogen but not triglycerides (Fig. 6a, b; Supplementary Fig. 9a-c). The opposing effect of obesity on the glycogen and triglyceride synthesis pathway was partially reflected in the induction of glycolytic gene expression in the obese mice liver but the impaired response of glycogen synthase to feeding-induced dephosphorylation and activation (Supplementary Fig. 9d, e). The induction of glucose-6-phosphate phosphatase in obesity may also contribute to the reduction in glycogen synthesis.

**Figure 6.**
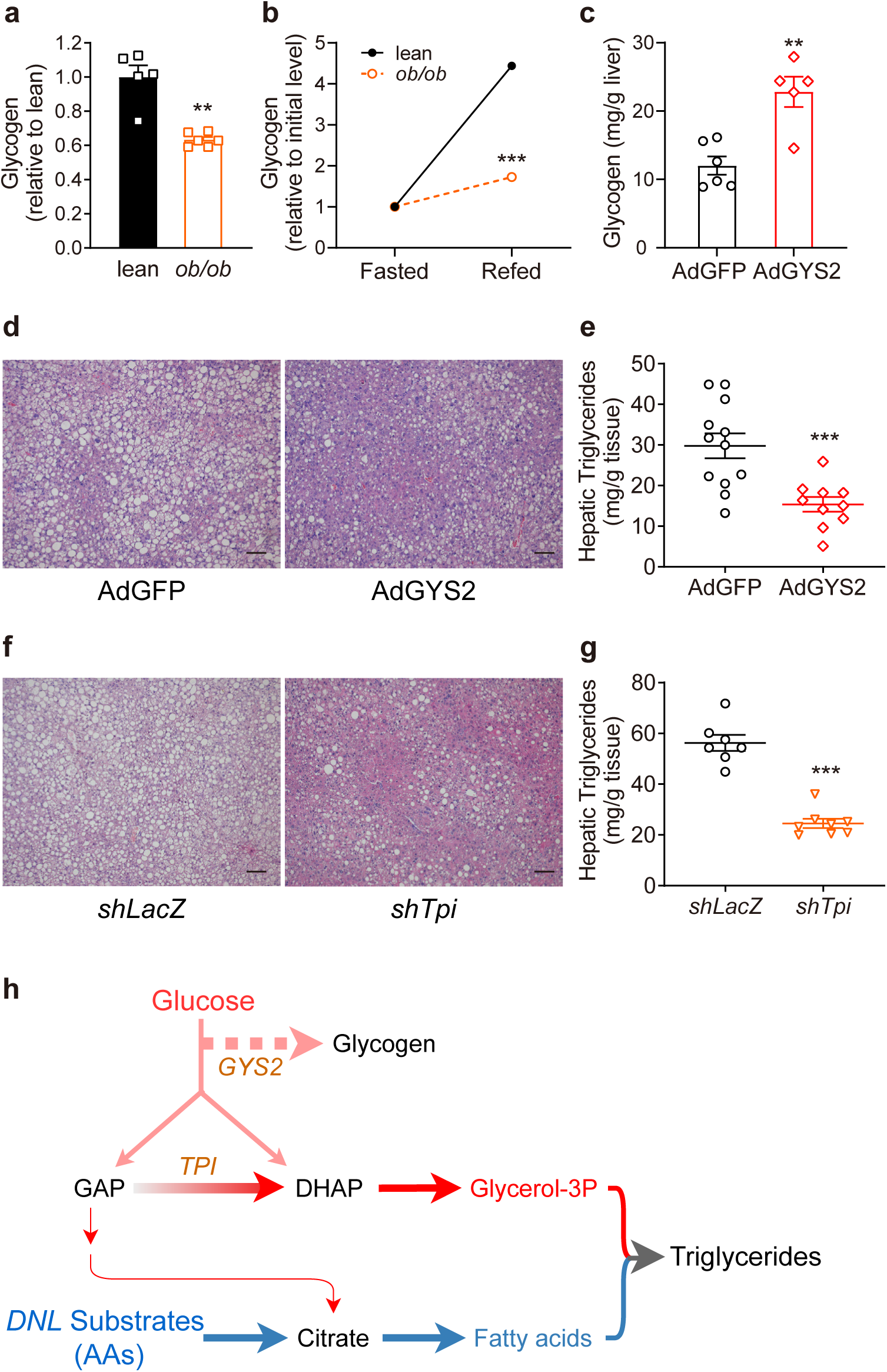
Inhibition of glycolytic glycerol-3P synthesis reduces hepatic triglyceride content. **a,** Glycogen quantification in lean (n=5) and *ob/ob* (n=6) mouse primary hepatocytes at basal state (4 hours after seeding). **b,** Measurement of glycogen synthesis in lean and *ob/ob* mouse primary hepatocytes. Hepatocytes were fasted in minimal medium for 2 hours and then incubated in complete medium (M199) containing 25 mM glucose, 100 nM insulin, 5% FBS for 4 hours (n=4). **c,** Measurement of hepatic glycogen levels in the *ob/ob* mice expressing either GFP (AdGFP, control, n=5) or GYS2 (AdGYS2, n=6), seven days post adenovirus administration. **d, e,** Haematoxylin & eosin (H&E) staining (**d**) and triglyceride measurement (**e**) of liver samples from *ob/ob* mice expressing GFP (n=12) and GYS2 (n=10). Scale bars, 50 μm. **f, g,** H&E staining (**f**) and triglyceride measurement (**g**) of liver tissues expressing shRNA targeting either *LacZ* (*shLacZ*, control, n=7) or *Tpi* (*shTpi*, n=8) in the *ob/ob* mice. Scale bars, 50 μm. **h,** Illustration of glycolytic flux contribution to lipid synthesis. All measurements are presented as means ± SEM; text labeled 0.05 < p < 0.1, *p < 0.05, **p < 0.01, ***p < 0.001, two-tailed unpaired Student’s *t-*test. N.D., not detected. See also Supplementary Figure 9.

To formerly test this hypothesis, we examined the effect of adenovirus-mediated glycogen synthase (GYS2) overexpression on hepatic TAG accumulation in the obese mice (Supplementary Fig. 9f). Consistent with our hypothesis, GYS2 overexpression not only doubled hepatic glycogen content (Fig. 6c) but also halved hepatic triglyceride content (Fig. 6d, e), without any significant effect on body weight as well as liver and adipose tissue size (Supplementary Fig. 9g). A similar beneficial effect of GYS2 overexpression on reducing hepatic triglyceride content was observed in the high-fat diet (HFD)-induced obese mice liver (Supplementary Fig. 9h), and the ability of the HFD obese mice to restore normoglycemia after a bolus injection of glucose was improved significantly (Supplementary Fig. 9i).

We then conducted a chemical inhibitor screening to further examine the specific contribution of individual glucose metabolism pathways toward hepatic TAG accumulation in obesity. As shown in Supplementary Fig. 9j, inhibition of glycerol-3P synthesis by GPDi significantly lowered triglyceride levels in the primary hepatocytes, whereas blocking the PPP (G6PDi) and glycolysis (PKMi) had no significant effect. Both *ex vivo* and *in vivo* flux analysis suggest hepatic glycerol-3P synthesis is synthesized by GA3P to DHAP conversion (Supplementary Fig. 9k). Therefore, we used adenovirus-expressed shRNA to specifically knockdown triosephosphate isomerase (TPI expression) in the obese mice liver (Supplementary Fig. 9l). As a result, hepatic triglyceride content was reduced by more than 60% (Fig. 6f-h), and no adverse effect on body weight and white adipose tissue size was observed (Supplementary Fig. 9m).

### Suppressing glutaminolysis driven reductive carboxylation or augmenting glutamine oxidation attenuates DNL

The intestinal absorption of glutamate and glutamine prevented us from directly assessing their contribution toward hepatic lipogenesis. Therefore, we resorted to chemical or genetic means to examine the contribution of the endogenous glutamate and glutamine pool toward hepatic triglyceride accumulation.

First, chemical inhibition of glutamine uptake by L-γ-glutamyl-p-nitroanilide (GPNA) had a similarly potent effect in reducing triglyceride accumulation in the primary hepatocytes as the inhibitor for fatty acid synthesis (ACCi, Fig. 7a). Then we performed broad suppression on glutaminolysis by liver-type glutaminase (GLS2) knockdown in the obese mice, and dramatically attenuated the accumulation of hepatic triglycerides (Supplementary Fig. 10a-c). Thus, these results confirmed that glutamine is a substantial source of the fatty acid synthesis in the liver.

**Figure 7.**
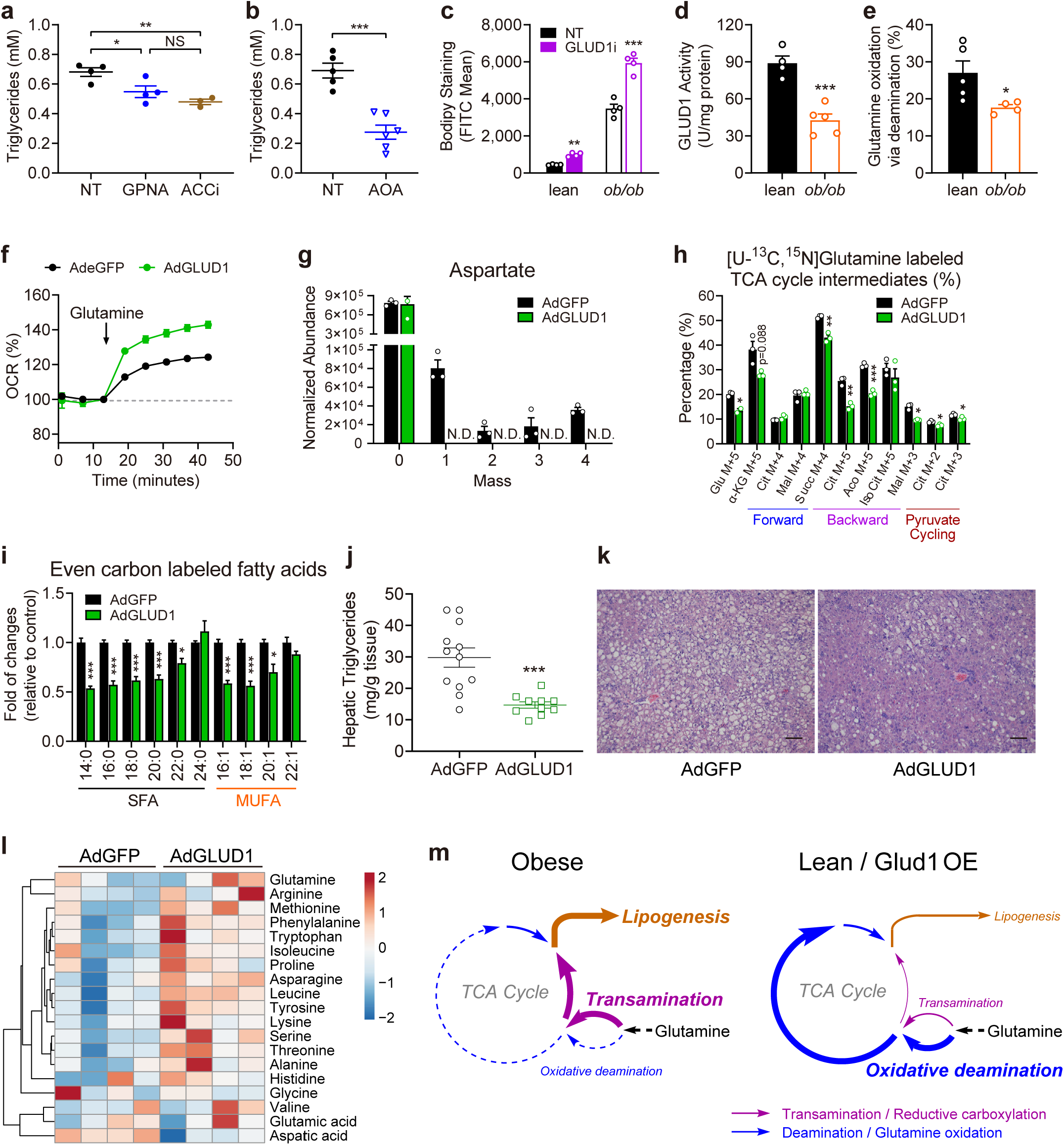
Broad and pathway-specific manipulation of glutamine metabolism alleviates hepatic steatosis. **a,b,** Triglyceride measurements of primary hepatocytes cultured in the presence or absence of glutamine transporter inhibitor L-γ-glutamyl-p- nitroanilide (GPNA, 2.5 mM, n=4, **a**), acetyl-CoA carboxylase inhibitor (ACCi, 10 μM, n=4, **a**), or transaminase inhibitor aminooxyacetate (AOA, 4 mM, n=6, **b**). **c,** Measurement of triglyceride content by BODIPY staining in primary hepatocytes cultured in the presence or absence of glutamate dehydroge-nase 1 inhibitor GLUD1i (R162, 40 μM, n=4) overnight. **d,** Measurement of GLUD1 activity in liver tissue samples prepared from lean and *ob/ob* mice (n=4 and 5 respectively). **e,** Measurement of glutamine oxidation capacity in primary hepatocytes prepared from lean and *ob/ob* mice as determined by the percentage of glutamine-induced oxygen consumption not suppressible by transaminase inhibitor AOA. n=3 for the lean, 4 for the *ob/ob* group. **f,** Measurement of glutamine oxidation in primary hepatocytes expressing either eGFP (control, n=4) or GLUD1 (n=3). (**g**) Isotopomer abundance of aspartate in [U-^13^C, ^15^N]Gln-traced (1 hour) primary hepatocytes prepared from *ob/ob* mice expressing either GFP (AdGFP, control) or GLUD1 (AdGLUD1), isolated three days post adenovirus administration, n=3. **h,** Fractional abundance of ^13^C-labeled TCA intermediates compared to the total (labeled and unlabeled) in [U-^13^C, ^15^N]Gln-traced (1 hour) primary hepatocytes treated as in (**g**). **i,** Fractional abundance of ^13^C-labeled triglyceride fatty acid (FA) species in primary hepatocytes treated as in (**g**). **j-k,** Triglyceride measurements (**j**) and H&E staining (**k**) of liver samples prepared from *ob/ob* mice expressing GFP (AdGFP, n=12, same control group as in Fig. 5g) or GLUD1 (AdGLUD1, n=10). Scale bars, 50 μm. **l,** Measurement of amino acid profiles in the liver tissue samples prepared from *ob/ob* mice expressing either GFP or GLUD1. Tissue samples were collected 10 days post virus administration. Color scale denotes row scaling z-scores. **m,** Schematic illustration of obesity-induced perturbation in hepatic glutamine metabolism. All measurements are presented as means ± SEM; text labeled 0.05 < p < 0.1, *p < 0.05, **p < 0.01, ***p < 0.001, two-tailed unpaired Student’s *t-*test. N.D., not detected. See also Supplementary Figure 10 and 11.

Our flux analysis and modeling studies revealed that the transamination and reductive carboxylation pathway of glutamate metabolism was increased in the obese primary hepatocytes while the oxidative deamination pathway might be suppressed (Fig. 2c, e, g; Fig. 3e; Supplementary Fig. 4g). Therefore, we hypothesized the reductive carboxylation pathway, not the oxidative deamination pathway, as the primary route of glutamate DNL. Consistent with this hypothesis, a general inhibition of transamination by AOA (Fig. 7b), or specific inhibition of glutamate-pyruvate transaminase (GPT) (Supplementary Fig. 11a), sharply reduced triglyceride accumulation, whereas treatment of primary hepatocytes with an inhibitor of the oxidative deamination enzyme, glutamate dehydrogenase (GLUD1), elevated cellular triglyceride content (Fig. 6c). Specific suppression of reductive carboxylation by adenovirus mediated knockdown of cytosolic isocitrate dehydrogenase 1 (IDH1) also significantly reduced hepatic triglyceride in obese mice liver (Supplementary Fig. 10a-c). We further confirmed the downregulation of the oxidative deamination pathway in the obese mice liver on both GLUD1 protein and its enzymatic activity levels (Fig. 7d, e). We therefore used adenovirus to overexpress GLUD1 (Supplementary Fig. 11b, c) as an effort to restore the glutamate oxidative deamination pathway and examine its impact on glutamine metabolism *ex vivo* and hepatic triglyceride accumulation *in vivo*.

As shown in Figure 7f, GLUD1 overexpression doubled the rate of glutamine oxidation, and it competitively suppressed the glutamate-driven transamination activity indicated by the abundance of ^15^N- labeled aspartate in obese primary hepatocytes (Fig. 7g). In addition, both the reductive carboxylation (backward) and pyruvate cycling were reduced significantly (Fig. 7h, Supplementary Fig. 11d, e). More importantly, fatty acid synthesis from glutamine was suppressed by GLUD1 overexpression (Fig. 7i). Intriguingly, incorporation of glucose carbon into fatty acids was also reduced to some extent in GLUD1- overexpressing primary hepatocytes (Supplementary Fig. 11f, g), probably through substrate competition in TCA cycle entrance or indirect contribution of glucose toward DNL through glutamine synthesis.

Consistent with flux analyses *ex vivo*, the overexpression of GLUD1 resulted in significant reduction in hepatic triglyceride accumulation in both the *ob/ob* and HFD obese mice models (Fig. 7j, k; Supplementary Fig. 9h, i). Glucose tolerance of the HFD obese mice was improved accordingly (Supplementary Fig. 11j). The suppression of DNL and improvement of hepatic steatosis occurred despite the elevation of the vast majority of amino acids except for glutamine and aspartate (Fig. 7l, m), confirming a prominent contribution from the endogenous glutamine pool in supporting DNL and the development of hepatic steatosis *in vivo*.

## Discussion

Historically, direct quantification of the carbon source for fatty acid synthesis has been difficult, and the estimates vary^17–20, 22, 34–38^. There are inherent challenges in the task, including heterogeneities within the different fatty acid pools, fatty acid turnover, and isotope dilution, combined with high abundance (1.1%) of background ^13^C in the nature. As has been demonstrated in a recent infusion study with 15 different ^13^C- labeled substrates from the Rabinowitz group, no labeling signals were detected in the circulating fatty acids^23^. There has been attempts to overcome this limitation by measuring ^13^C-enrichment of tissue acetyl-CoA or malonyl-CoA pool as a surrogate for DNL^20, 29^. Current LC-MS technology has been mature enough to measure acetyl-CoA and malonyl-CoA at low nanomole/mL levels^39, 40^. However, due to the complexity of metabolic pathways and compartmentalization of acetyl-CoA and malonyl-CoA, a quantitative relationship between ^13^C-enrichment acetyl-CoA or malonyl-CoA pool with DNL has to be established before meaningful interpretations can be made.

Nevertheless, the contribution of glucose and glutamine toward the TCA cycle is very similar in our primary hepatocyte studies presented herein as those reported by mouse infusion studies^23^, suggesting primary hepatocytes are an invaluable system for *in vivo* approximation and hypothesis generation. We were further able to confirm those findings *in vivo* through bulk feeding and genetic models. Through a combined approach, we were able to not only account for over 70% of all hepatic DNL carbon sources for the first time but also demonstrate deregulations in glycogen synthesis and reductive carboxylation pathways as the primary cause of hepatic steatosis in obesity.

Additionally, by comparing *ex vivo* and *in vivo* flux studies, we observed intestinal absorption as a major modifier of dietary metabolism. Under *ex vivo* conditions, glucose carbon primarily fills the glycolysis, pentose phosphate pathway, and glycerol-3-phosphate synthesis pathway, while TCA and its surrounding pathways are dominated by amino acids (Fig. 1l, 3b). Only under nutrient imbalanced conditions, such a territorial barrier in metabolism is lost, and glucose and amino acid carbon backbone may enter each other’s metabolic orbit (Supplementary Table 1). Such a castling and complementation phenomenon in carbohydrate and amino acid metabolism reduces futile cycle under nutrient rich conditions while enables metabolic flexibility under nutrient imbalanced/restricted conditions.

However, under *in vivo* conditions, the territorial metabolism of glucose and amino acids are overrun by extra-hepatic transformation of glucose into organic acids, which could be readily taken up by the liver to fuel the TCA cycle and enter the DNL pathway. Besides glucose, intestinally absorbed amino acids are also transformed into organic acids in circulation, which may facilitate extra-hepatic utilization of amino acids for energy production, lipogenesis, and nucleotide biosynthesis. Additionally, amino acids cleared by the intestine may feed the microbiota to further modulate host nutrient metabolism and lipogenesis. Further, nutrients may participate in metabolism not only as a substrate but also a transcriptional/allosteric regulator. At least in the case of fructose, hepatic fructolysis is necessary to drive lipogenic gene expression and the development of metabolic syndrome^41, 42^, and fructose-to-organic acid transformation in the intestine reduce hepatic steatosis^43^, indicating the importance of nutrient both as a signal and a substrate in driving lipogenesis. Therefore, further genetic and combinatory flux analyses *in vivo* and *ex vivo* are critically needed to untangle tissue crosstalk in nutrient metabolism and evaluate the role of nutrient balance in physiology and obesity.

## Methods

### Mouse Husbandry and Experimentation

All the animal husbandry and experimental procedures were in strict accordance with the guidelines approved by the Institutional Animal Care and Use Committee (IACUC) of Tsinghua University. Male leptin-deficient (*ob/ob*) mice and lean wild-type littermates of 8-12 week old were obtained from HFK Bioscience (Beijing, China). For the diet-induced obesity model, male C57BL/6J mice purchased from Charles River (Beijing, China) were placed on a 6-month high-fat diet (D12492: 60% kcal% fat; Research Diets) starting from 4 weeks after birth. All the mice were housed in a specific pathogen free (SPF) facility at Tsinghua Laboratory Animal Research Center under 12/12-hours light-dark cycle and controlled temperature (25 ± 1°C) with free access to water and food.

### Primary Hepatocyte Isolation and Culture

For primary hepatocyte isolation, lean and *ob/ob* mice from 8-12 weeks old were first anesthetized with 80 mg kg^-1^ sodium pentobarbital. The mouse liver was first perfused with 40mL of warm (37°C) HBSS solution (Corning, Cat#21-022-CVR) containing 1mM EGTA (Amresco, Cat#0732) and 5.5mM glucose at a speed of 11 mL min^-1^, and then digested with 50 (lean) or 75 (*ob/ob*) mL of 0.3mg/mL Collagenase Type IV (Sigma, Cat#C5138) solution prepared in HBSS supplemented with 5mM CaCl_2_ (Solarbio, Cat#C8370) and 5.5mM glucose. Primary hepatocytes were dispersed and sedimented at 50g for 2 (lean) or 7 (*ob/ob*) minutes, washed twice with M199 (MacGene, Cat#CM10013), and re-suspended in the attachment medium (M199 with 0.2% BSA, 2% FBS (Excell, Cat#11H116), 1% penicillin/streptomycin). Live cell numbers were determined by Trypan Blue (Amresco, Cat#K940) exclusion. After cell counting and normalization, primary hepatocytes were seeded onto 0.1% gelatin (Amresco, Cat#9704) coated culture plates, incubated at 37°C with 5% CO_2_ for 4-8 hours, washed, and maintained in M199 with 0.1mg/mL Primocin (InvivoGen, Cat#ant-pm-2) until further experimentation. Primary hepatocytes from mice maintained on normal chow diet (NCD) and HFD (high-fed diet) were isolated and traced with the same procedures.

### *Ex vivo* ^13^C-Isotopomer Tracing

Mouse primary hepatocytes were seeded in 100 or 60mM plates at a density of 3-3.5×10^6^ or 1.6 ×10^6^ cells per plate, respectively. After 6 hours, hepatocytes were washed with phosphate buffer solution (PBS) and pre-incubated in a base medium without glucose or amino acids for 45 minutes to deplete the intracellular pool of unlabeled macronutrients. The primary hepatocytes were then replaced in a fresh base medium supplemented with the isotopic tracers, 10mM [U-^13^C]glucose (CLM, Cat#CLM-1396-PK) or 2.5mM [U- ^13^C,^15^N]glutamine (CLM, Cat# CNLM-1275-H-PK). At indicated time points, media were promptly drained and plates were snap-frozen in liquid nitrogen. To label newly synthesized fatty acids, primary hepatocytes were first pre-incubated in M199 supplemented with 100nM insulin (Sigma, Cat#10516) for lipogenesis activation, and incubated in the same media containing either 15mM (5mM ^12^C+10mM ^13^C) glucose plus 3.2mM ^12^C glutamine (Gibco, Cat#25030-081), or 15mM ^12^C Glucose plus 3.2mM (0.7mM ^12^C,^14^N+ 2.5mM ^13^C,^15^N) glutamine, or 15mM ^12^C glucose plus 0.365 g/L [U-^13^C,^15^N]amino acids mixture (CLM, Cat#CNLM-452) for 6 hours. M199 contains 50mg/L of sodium acetate.

### *In vivo* ^13^C-Isotopomer Tracing

To assess the contribution from dietary carbohydrates and amino acids toward *in vivo* energy metabolism and lipogenesis, mice were fed for 24 hours with glucose and amino acids dissolved in drinking water at a 3:1 ratio (15% glucose + 5% amino acids mixture *w/w*) mimicking normal chow diet formulation (LabDiet 5K52). The drinking water contained either [U-^13^C]glucose (CLM, Cat#CLM-1396) plus unlabeled ^12^C^14^N amino acids (ULM-7891), or ^12^C glucose plus [U-^13^C, ^15^N]amino acids (CNLM-6696). Plasma and tissues were harvested with liquid nitrogen. For metabolites extraction, serum (80μl) or liver tissue (50mg) was homogenized in 2mL pre-chilled (-80°C) 80% (*v/v*) methanol solvent. For triglyceride fatty acids extraction, liver tissue (50mg) was homogenized in 3mL pre-chilled (-20°C) 50% (*v/v*) methanol solution with 0.1M HCl. The samples were processed as the following extraction steps, and used for LC–MS analysis.

### Metabolite Extraction and Metabolomics Analysis

To extract metabolites from primary hepatocytes, 2 or 4mL of pre-chilled (-80°C) 80% (*v/v*) methanol solvent was added to each 60mm or 100mm plate respectively, and the extraction was placed at -80°C for 20 min. Cells were then scraped down from the plate on dry ice and transferred to an Eppendorf tube. For tissue sample extraction, frozen liver tissues were weighed (∼50mg) and homogenized in 20 volumes of the same methanol solvent using polytron homogenizer (32000rpm for 30 seconds). The lysates were cleared by centrifugation at 14000g (4°C) for 5 minutes. The supernatant was then transferred to a new tube, air-dried in a Speed-Vac and stored at -80°C until mass spectrometry analysis.

The dried extracts were re-dissolved and analyzed using a targeted and untargeted liquid chromatography mass spectrometry (LC-MS/MS). Glycolysis and TCA cycle metabolites were measured on a TSQ Quantiva Triple Quadrupole mass spectrometer (Thermo Fisher Scientific, CA) with positive/negative ion switching. For amino acids profiling, a Q Exactive orbitrap mass spectrometer (QE-MS, Thermo Fisher Scientific, CA) was applied. The integrated peak intensities obtained from LC-MS were used for further data analysis.

For metabolic flux analysis, isotope labeling was corrected for natural ^13^C abundance. Between-group differences of metabolites abundances were assessed by Welch’s two-sample t-test, with p < 0.05 considered as statistically significant. For untargeted metabolomics analysis, raw values of each sample were normalized by its median of total metabolites intensity. Any missing values were assumed to be below the detection limit and were imputed with half of minimum of the whole dataset. Then hierarchical clustering (HCL) and principal components analysis (PCA) were employed to determine the data quality and detect potential outliers. Metabolites with more than two-fold change in normalized values and less than 0.05 in p-values were selected as differentially regulated. The statistical analysis and heat map visualization were performed using R scripts (http://cran.r-project.org/) executed in RStudio (v. 0.99.896) for Windows.

### Total Fatty Acids Extraction and Flux Analysis

For total lipid extraction, 50% (*v/v*) methanol solution with 0.1M HCl was prepared and pre-chilled at - 20°C. The primary hepatocytes in 100mM dishes were scraped down using the methanol solution and transferred into 10mL glass tubes. An aliquot of 1.5mL chloroform was added into each tube and vigorously mixed with vortex for 1 minutes. The samples were centrifuge at 3000 rpm for 15 minutes to achieve phase separation, and 1mL of the bottom layer was carefully transferred to a new glass tube, and evaporated by a stream of nitrogen. Fatty acids from total lipids were released by saponification (refluxing with 3mL 0.3M KOH in 90% methanol at 80°C for 1 hour and remaining ∼300μL solvent). The samples were acidized with 300μL formic acid (Fisher Scientific, Cat#A117-50), and then extracted with 3mL N-hexane. After a brief vortex, the mixture was allowed to settle naturally, and the top layer was transferred to a new glass tube. The lower layer were reextracted with another 1mL N-hexane, combined and air-dried under a constant stream of nitrogen. The dried samples were directly sent for LC-MS analysis.

### Measurements of Triglycerides and Glycerol

Total lipids from liver tissues (∼100mg) were extracted with chloroform/methanol (2:1, *v/v*) according to the Folch method^44^ and air-dried in a Speed-Vac. The samples were re-solubilized in 5% NP-40 with two cycles of 95°C, 5 minutes. Total lipids from primary hepatocytes (∼0.2 million) were directly extracted with 5% NP-40 (MP, Cat#0219859680). Triglyceride levels were measured by triglyceride assay kit (Nanjing Jiancheng Bioengineering Institute, Cat#A110-2-1). Medium or intracellular glycerol levels were measured by glycerol assay kit (Applygen, Cat#E1002).

### Reverse Transcribed Quantitative Polymerase Chain Reaction (RT-qPCR)

Total RNA was extracted from liver tissues with TRIzol reagent (Life Technologies, Cat#15596018) according to manufacturer’s recommendations. An aliquot of 0.5 μg RNA were reverse transcribed to cDNA by FastKing RT Kit (KR116-02, Tiangen). cDNA samples were used as templates to perform real-time quantitative PCR with the power SYBR green mix (Abm, Cat#MasterMix-LR) on an Applied Biosystems QuantStudio (TM) 7 Flex System (Applied Biosystems). Duplicate runs of each sample were normalized to *36b4* to determine the relative transcript abundance of target genes. RT-qPCR primer sequences as following (F, forward; R: reverse):

*36b4*: F: 5’-CTTCATTGTGGGAGCAGACA-3’, R: 5’-TCTCCAGAGCTGGGTTGTTC-3’;

*Tpi*: F: 5’-CCAGGAAGTTCTTCGTTGGGG-3’, R: 5’-CAAAGTCGATGTAAGCGGTGG-3’;

*Gpd1*: F: 5’-ATGGCTGGCAAGAAAGTCTG-3’, R: 5’-CGTGCTGAGTGTTGATGATCT-3’;

*Acss2*: F: 5’-CCAGGAAGTTCTTCGTTGGGG-3’, R: 5’-CAAAGTCGATGTAAGCGGTGG-3’.

### Protein Extraction and Western Blot Analysis

Total protein lysates were extracted from liver tissues or cells with ice-cold radioimmuno-precipitation assay (RIPA) lysis buffer (50mM Tris pH 7.5, 150mM NaCl, 0.1% SDS, 1% NP-40, 0.5% sodium deoxycholate) supplemented with protease and phosphatase inhibitor cocktails (Pierce, Cat#78441). The lysates were clarified by centrifugation at 13000 rpm for 15 minutes, and protein concentrations were quantified by BCA (Solarbio, Cat#PC0021). Equal amounts (50μg) of protein lysates were resolved by SDS-PAGE, blotted onto a nitrocellulose membrane (Pall, Cat#66485), blocked with 5% milk in TBST for 1 hour at room temperature, then incubated overnight with primary antibodies at 4°C. The blots were then washed and incubated with HRP-conjugated secondary antibody for 1 hour at room temperature, and visualized by ECL (Merck Milipore, Cat#WBKL50500) using a ChemiDoc MP imaging system (BioRad). For the detection of phosphoepitopes, BSA instead of milk was used for blocking and antibody preparation. Antibodies used in this study are listed below.

Rabbit anti-Glycogen Synthase (Cell Signaling Technology, Cat#3886S)

Rabbit anti-phospho-Glycogen Synthase (Ser641) (Cell Signaling Technology, Cat#3891)

### GLUD1 (Santa Cruz, Cat# sc-160382)

Rabbit anti-ACL (CST, Cat#4332)

Rabbit anti-ACSL1 (CST, Cat#9189)

Rabbit anti-FASN (CST, Cat#3180)

Mouse anti-HIF1A (Abcam, Cat#ab16066)

Rabbit anti-PDH (CST, Cat#3205)

Rabbit anti-GCK (PTG, Cat#19666-1-AP)

Rabbit anti-PCK1 (PTG, Cat#16754-1-AP)

Mouse anti-β-Actin (Abgent, Cat#AM1021B)

Mouse anti-GAPDH (Easybio, Cat#BE0023)

### Measurement of Mitochondria Respiration and Glycolysis

Mitochondria respiration and glycolysis were measured with the Seahorse XF96^e^ extracellular flux analysis system (Agilent). Fresh primary hepatocytes were plated onto gelatin-coated, XF 96-well microplates at a density of 4×10^3^/well. The medium was replaced after 4-8 hours with XF assay medium (Seahorse Biosciences 103334-100, supplemented with 10mM glucose, 2mM pyruvate, 4mM glutamine) or minimal medium containing same concentration of salt ions. The oxygen consumption rate (OCR) and extracellular acidification rate (ECAR) were measured by XF^e^96 Extracellular Flux Analyzer (Seahorse Biosciences) according to manufacturer’s recommendations. Reagents were used at the following final concentrations: glucose/fructose (10mM), pyruvate (2mM), glutamine (4mM), oligomycin (4μM), FCCP (0.5μM), rotenone (2μM), antimycin A (2μM). For the mitochondria fuel test, inhibitors were used at the following concentrations: DMSO (0.1%), etomoxir (40μM, MCE, Cat#HY-50202), UK5099 (40μM, MCE, Cat#HY- 15475), aminooxyacetate (AOA, 40mM, Macklin, Cat#0805087). Measured values were normalized to total protein amount quantified by BCA or cell number of the primary hepatocytes plated on parallel plates.

### Adenovirus-mediated Knockdown and Overexpression

For knockdown experiments, adenoviruses expressing *LacZ* and *Tpi* shRNA were cloned into the Adenoviral Gateway Expression System (Invitrogen, CA). In brief, the shRNA oligos targeting *LacZ* and mouse *Tpi* obtained from MISSION shRNA Library (Sigma) were first cloned into their respective entry vectors (pENTR-U6), and then recloned into their corresponding destination vectors (pAd/BLOCK-iT™- DEST) through LR recombination. For overexpression experiments, PCR amplification of *Gfp* (control) and *Glud1* open reading frame from mouse cDNA were first cloned into pENTR1A and then recombined into pAd/CMV/V5-DEST Gateway®.

Primer sequence used for cloning shRNA targeting *Tpi*: top-strand (5’-3’):

CACCGTTCGAGCAAACCAAGGTCATCTCCGAAGAGATGACCTTGGTTTGCTCGAAC,

bottom-strand (5’-3’):

AAAAGTTCGAGCAAACCAAGGTCATCTCTTCGGAGATGACCTTGGTTTGCTCGAAC.

Primer sequence used for cloning shRNA targeting *Acss2*:

top-strand (5’-3’): CACCGCGAATGCCTCTACTGCTTTCTCGAGAAAGCAGTAGAGGCATTCGC,

bottom-strand (5’-3’): AAAAGCGAATGCCTCTACTGCTTTCTCGAGAAAGCAGTAGAGGCATTCGC.

For virus packaging and expression, plasmids were first digested and then transfected into HEK293A cells. After three rounds of amplifications, virus-producing HEK293A cells were harvested and lysed, and the released virus particles were purified by 55% cesium chloride and desalted through a PD10 column (GE Healthcare Life Sciences, Cat#17085101). Adenovirus particles were tail vein injected at doses of 0.1 OD for the lean, wild-type mice, or 0.12-0.15 OD per *ob/ob* mice. Mice were sacrificed 7-10 days after virus administration following a 6-hour food withdrawal, and the collected tissues were either immediately fixed in a 4% (*v/v*) paraformaldehyde (PFA) or snap-frozen in liquid nitrogen and stored in -80°C. For *ex vivo* overexpression studies, primary hepatocytes were infected with adenoviruses at a concentration of 6.25× 10^-4^ OD/1×10^5^ cells for 24 hours.

### Histology Analysis

For hematoxylin and eosin (H&E) staining, liver specimens were promptly fixed in 4% PFA after harvesting, dehydrated, and paraffin-embedded. The embedded tissues were then sectioned into 5-μm-thick sections and stained with hematoxylin/eosin according to the standard protocols.

### Glucose and Insulin Tolerance Tests

On day 7 and 9 after adenovirus injection, glucose tolerance tests (GTT) were performed in overnight-fasted HFD mice by intraperitoneal glucose injection (2 g kg^−1^ body weight), and insulin tolerance tests (ITT) were performed by intraperitoneal insulin injection (1.5 IU kg^−1^ for HFD mice) after a 6 hours food withdrawal. Tail plasma glucose concentrations were measured before and after the injection at the indicated time points.

### Cell Treatment and Lipid Droplet Staining

Primary hepatocytes were incubated with different substrates or inhibitors for 16-24 hours at following concentrations: 10mM glucose (Amresco, Cat#0188-500G); 4mM glutamine (Gibco, Cat#25030-081); inhibitors targeting ACC (10nM MK-4074, MCE, Cat#HY-107709), GPD (10μM ginkgolic acid 15:1, Push Bio-technology, Cat#PB0667-0005), G6PD (1M 6-aminonicotinamide, Sigma, Cat#329-89-5), PKM (20μM Compound 3K, MCE, Cat#HY-103617), SLC1A5 (2.5mM L-γ-glutamyl-p-nitroanilide, GPNA, Aladdin, Cat#S161136), GLUD1 (40μM R162, MCE, Cat#HY-103096). Then the cells were washed with PBS and fixed with 4% PFA for 15min. The lipid droplets were then stained with 0.38μM BODIPY 493/503 fluorescent dye (Thermo Fisher Scientific, Cat#D3922) at 37°C for 5min and imaged. For flow cytometry-based quantification of lipid droplets, cells were counter-stained with DAPI (Solarbio, Cat#C0060) before being released from culture dish by trypsinization (Macgene, Cat#CC017). A total of 10,000 events were analyzed per sample. Average BODIPY signals of live cells were used for subsequent analyses.

### Glycogen Measurement

Primary hepatocytes glycogen contents were measured by glycogen assay kit (Bioassay, Cat#E2GN-100). Alternatively, samples were mixed with aqueous alkali, and incubated at 100°C for 15 minutes to solubilize glycogen and quantified following the manufacture’s recommendations (Nanjing Jiancheng Bioengineering Institute, Cat#A043-1-1).

### Glutamate Dehydrogenase Activity Assay

Liver tissues (∼100mg) were homogenized in PBS, and the glutamate dehydrogenase (GLUD1) activity was directly measured by α-ketoglutarate-dependent NAD^+^ formation in the presence of NH_4_ and NADH according to manufacturer’s recommendation (Solarbio, Cat#BC1460).

### Glutamine and Glutamic acid Determination Assay

Glutamine and glutamic acid of liver tissues (∼50mg) or primary hepatocytes (1.6×10^6^ cells) were extracted by pre-chilled (-80°C) 80% (*v/v*) methanol solvent following the metabolite extraction method. Samples were air-dried and re-dissolved in acetonitrile. The concentration of glutamine in the samples was then determined by LC-MS. The concentration of glutamic acid in the sample was determined chemically according to manufacturer’s recommendations (Nanjing Jiancheng Bioengineering Institute, Cat#A074-1- 1), and the relative concentrations were confirmed by LC-MS.

### Metabolic Flux Analysis

Mass isotopomer distributions of the detected intracellular metabolites, citrate, α-ketoglutarate, malate (as a combined node of succinate/succinyl-CoA/fumarate/malate), lactate and glutamic acid were used for the fluxes estimation. A simplified TCA cycle network based on a framework from reductive carboxylation research^45^ was applied for calculation, and the flux analysis was performed by OpenMebius^25^. Isotopically nonstationary ^13^C-MFA (INST-MFA)^46^ method was employed for glucose tracing results with multiple time points (5, 30, 120 minutes), and conventional ^13^C-MFA^47^ were used for analyzing glutamine tracing results, for its rapid plateau at 10 minutes. The initial metabolite pools in the network were set as the intraheptocellular concentrations collected by HepatoDyn^48^.

### Mass Isotopomer Enrichment-based Estimate of DNL

*D* represents the fractional contribution of ^13^C labeled substrate carbon to lipogenic acetyl-CoA (AcCoA), and *S* represents the fraction of newly synthesized fatty acid among the total pool for any given fatty acid species (e.g., palmitate). Mass isotopomer enrichment (MIE) and mole percent enrichment (MPE) represents the percentile enrichment of palmitate isotopomers and ^13^C atoms, respectively, for the same fatty acid.

In the case of palmitate, it has eight potential ^13^C-containing isotopomers (MIE_1∼8_): M+2, M+4, M+6, M+8, M+10, M+12, M+14, M+16. All odd number isotopomers are excluded due to their lower abundance. If we use MIE_i_ to denote the percentile abundance of the palmitate isotopomer that has *i* ^13^C_2_ units, then:

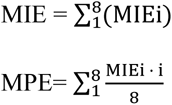

The solution for *D* and *S* for palmitate can be found by solving the following two equations in Matlab:

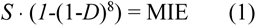

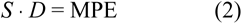

*D* and *S* can also be estimated by the Fatty Acid Source Analysis (FASA) program in Matlab^31^, which is based on the principle of mass isotopomer distribution (MID).

### Statistical Analysis

Data are presented as mean ± SEM. Student’s unpaired t-test was performed to determine statistical differences between two groups. Multiple groups or time-series comparison for substrate flux analysis and GTT were assessed by two-way ANOVA. *P*-values of less than 0.05 were considered as significant, and *p*- values ranging from 0.05 to 0.1 were labeled on the graphs of metabolic flux. N denotes the number of biological replicates in each experiment, and it is provided in the corresponding figure legends. Immunoblot quantification was completed by AlphaView (FluroChem FC3). Schematic diagrams and cartoons were drawn in ChemDraw 17.0 or Adobe Illustrator CS6. All other graphs were drawn in Graphpad Prism 8.0.2 software.

## Acknowledgements

We thank X. Liu, X. Wang, Y. Jiao, and L. Xu from the Metabolomics Facility at Technology Center for Protein Sciences and W. Wang from the Pharmaceutical Technology Center at Tsinghua University for technical help about LC-MS and metabolomics data analysis. We thank the Core Facility of Center for Biomedical Analysis, Tsinghua University for technical support about Seahorse analysis. We thank Peng Li, Guangshuo Ou, Yiguo Wang, Xiaowei Chen, Ligong Chen, and Peng Jiang for helpful discussions with this project. We thank H. Huang, N. Li and all present and former membranes of the Fu lab for their technical assistance and discussions. This work is supported by the National Science and Technology Major Project (2016YFA0502002 and 2017YFA0504603), and National Natural Science Foundation of China (NSFC 81471072 and 31671229). Additional support for the Fu lab comes from Tsinghua-Peking Center for Life Sciences, National 1000 Junior Scholar Program, DOE Key Laboratory of Bioinformatics, and the Center for Synthetic and Systems Biology at Tsinghua University.

## Author Contributions

Conceptualization: Y.L and S.F; Investigation: Y.L, L.L, H.L, X.B, F.S; Writing: Y.L and S.F; Supervision: S.F.

## Competing Interests Statement

The authors declare no competing financial interests.

---

**Supplementary Figure 1.**
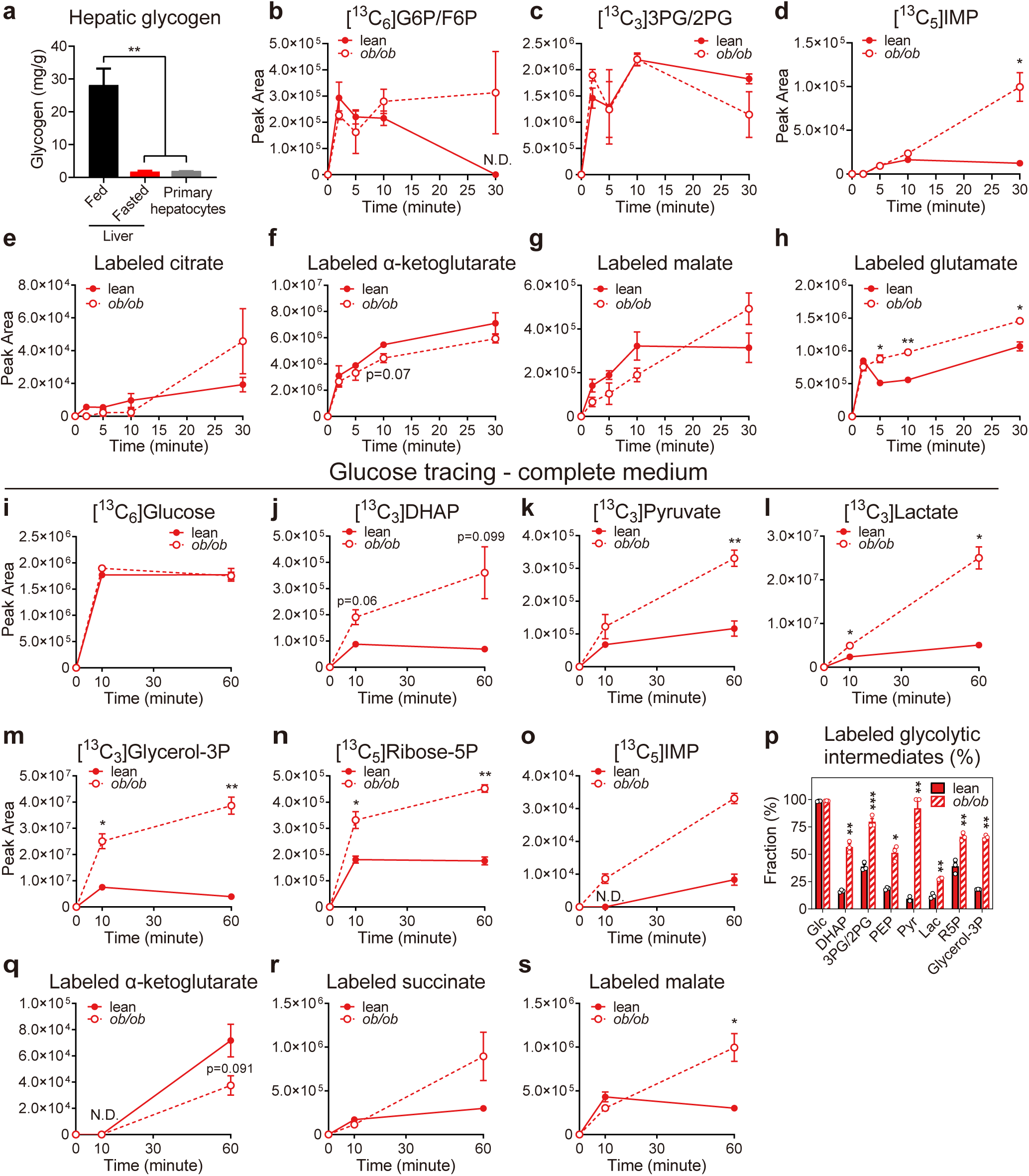
Related to Figure 1; Other glycolysis intermediates, and the glucose flux in complete medium a, Glycogen level of liver from fed and overnight fasted mice compared with that of primary hepacytes ready for tracing (4 hours after isolation). **b-d,** Accumulation of uniform,^13^C isotope-labeled glucose and metabolites in lean and *ob/ob* mice primary hepatocytes during the course of [U-^13^C]Glc tracing in minimal medium. Glucose, Glc; glucose 6-phosphate/fructose 6-phosphate, G6P/F6P; 3-phosphoglycerate/2-phospho- glycerate, 3PG/2PG; inosine monophosphate, IMP. **e-h,** Accumulation kinetics of ^13^C-labeled TCA cycle intermediates in primary hepatocytes traced in minimal medium. **i-o,** Accumulation kinetics of uniform,^13^C isotope-labeled glucose and metabolites in primary hepatocytes traced in complete medium (DMEM containing 4 mM glutamine and other amino acids). **p,** Fractional abundance of [U-^13^C]-labeled glycolysis intermediates in hepatocytes traced in complete medium compared to the total of corresponding metabolites at steady state (60 minutes). **q-s,** Accumulation kinetics of ^13^C-labeled TCA cycle intermediates in hepatocytes traced in complete medium.

**Supplementary Figure 2.**
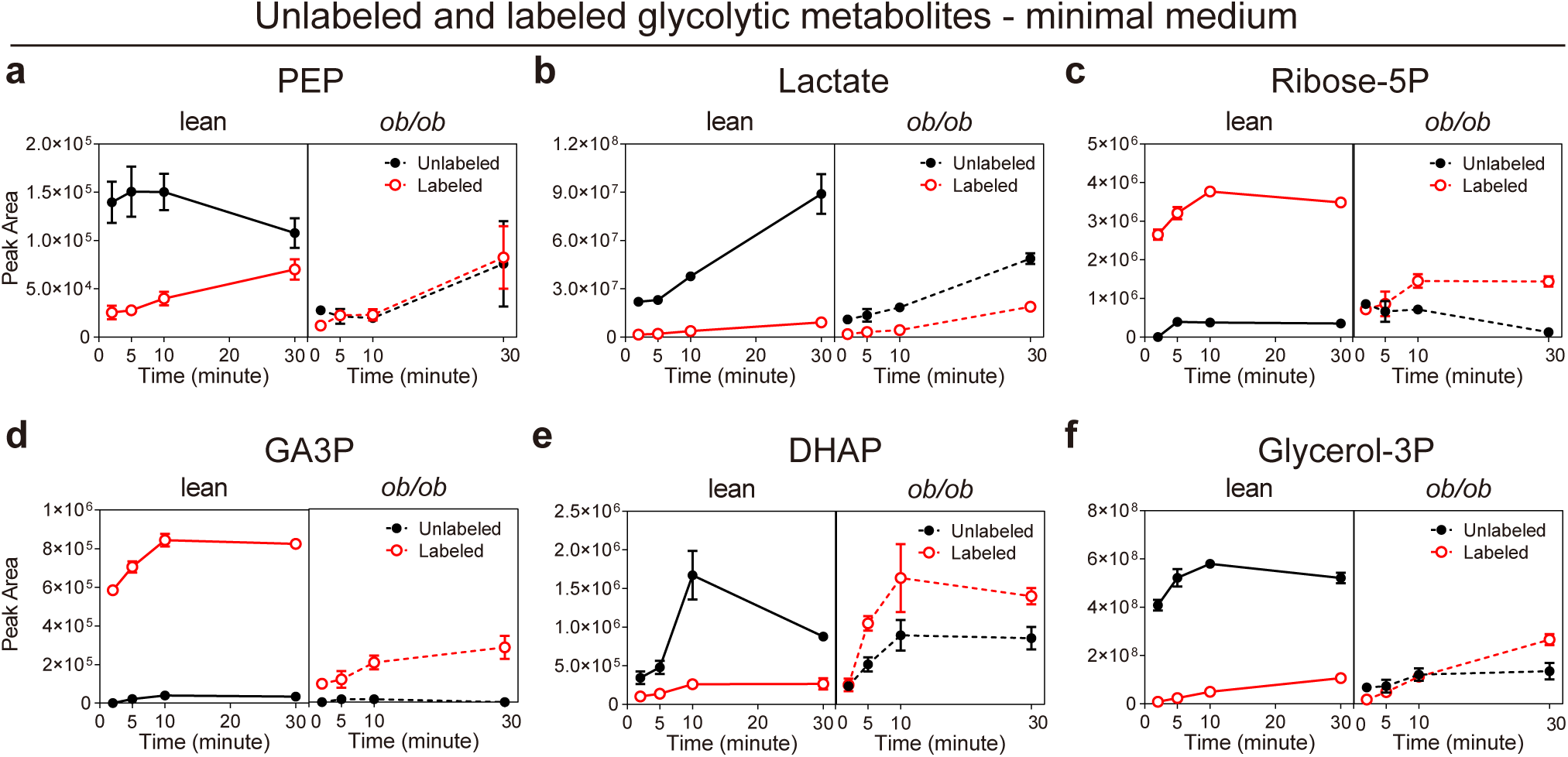
Related to Figure 1; Comparison between glucose and non-glucose substrates flux a-f, Changes of unlabeled and ^13^C-labeled TCA cycle intermediates abundance in hepatocytes traced in minimal medium, the same experiment with Fig. 1. Data are presented as means ± SEM, n=3.

**Supplementary Figure 3.**
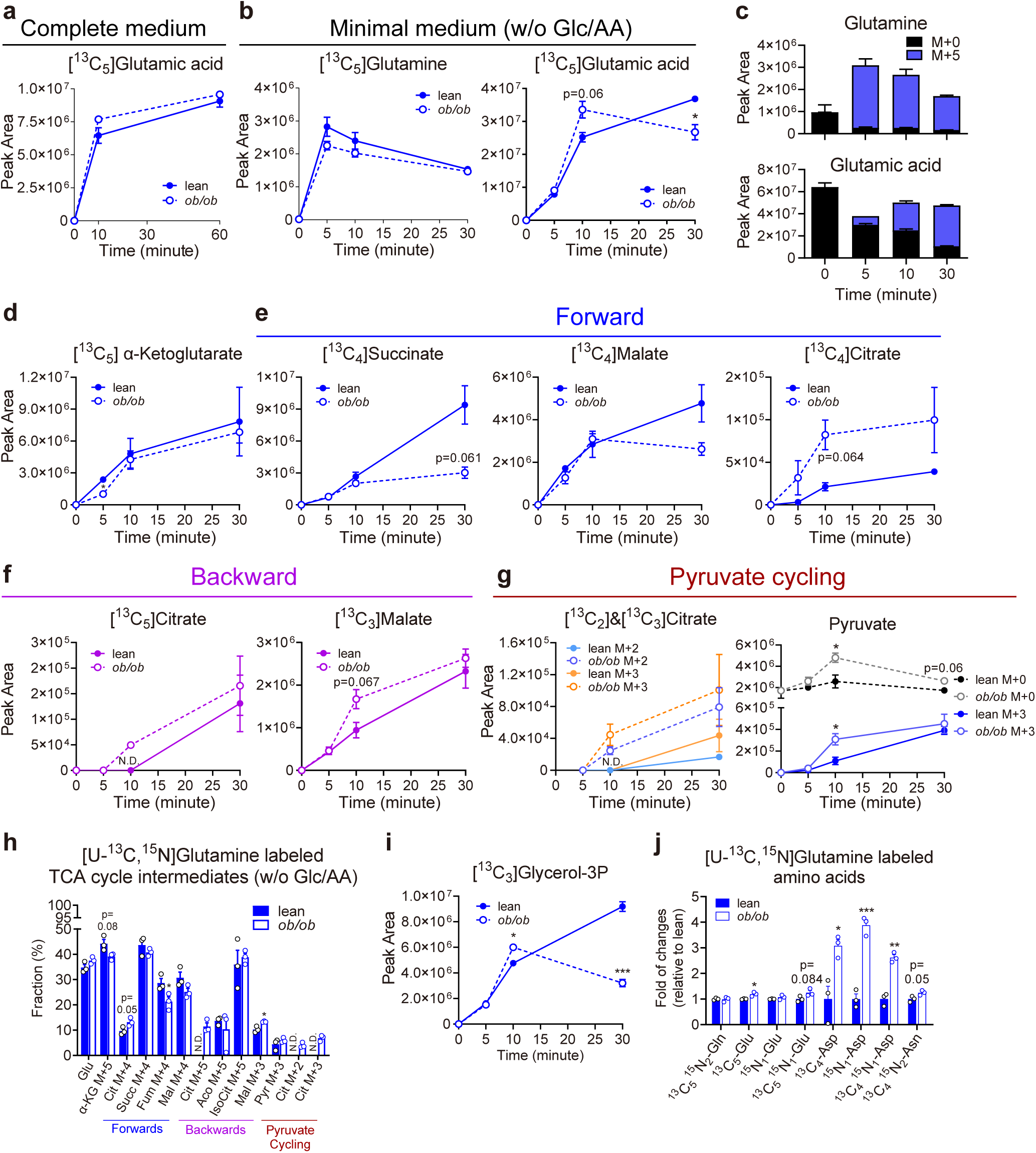
Related to Figure 2; Glutamine flux of lean and *ob/ob* mice primary hepatocytes without other substrates. **a,** Accumulation kinetics of ^13^C isotope-labeled glutamic acid (M+5) in lean and *ob/ob* mice primary hepatocytes after the addition of [U-^13^C, ^15^N]glutamine in complete medium. **b,** Accumulation kinetics of ^13^C-labeled glutamine (left) and glutamic acid (right) isotopomers (M+5) in hepatocytes traced in minimal medium. **c,** Abundance of glutamine (upper) and glutamic acid (lower) M+0 or M+5 isotopomer in hepatocytes during tracing in minimal medium. **d-g,** Accumulation kinetics of ^13^C isotope-labeled TCA cycle intermediates in hepatocytes traced in minimal medium. **h,** Accumulation kinetics of ^13^C isotope-labeled glycerol-3P (M+3) in lean and *ob/ob* mice primary hepatocytes during [U-^13^C, ^15^N]Gln tracing. **i,** Fractional abundance of all ^13^C isotope-labeled TCA cycle intermediates compared to their respective total in the primary hepatocytes 10 minutes after the addition of by [U-^13^C, ^15^N]Gln in minimal medium. Glutamate, Glu; oxaloacetate, OAA; citrate, Cit; aconitate, Aco; isocitrate, IsoCit; α-ketoglutarate, α-KG; succinate, Succ; malate, Mal. **j,** Fractional abundance of ^13^C or ^15^N-labeled amino acids in lean mice primary hepatocytes at 10 minutes after the addition of [U-^13^C, ^15^N]Gln.

**Supplementary Figure 4.**
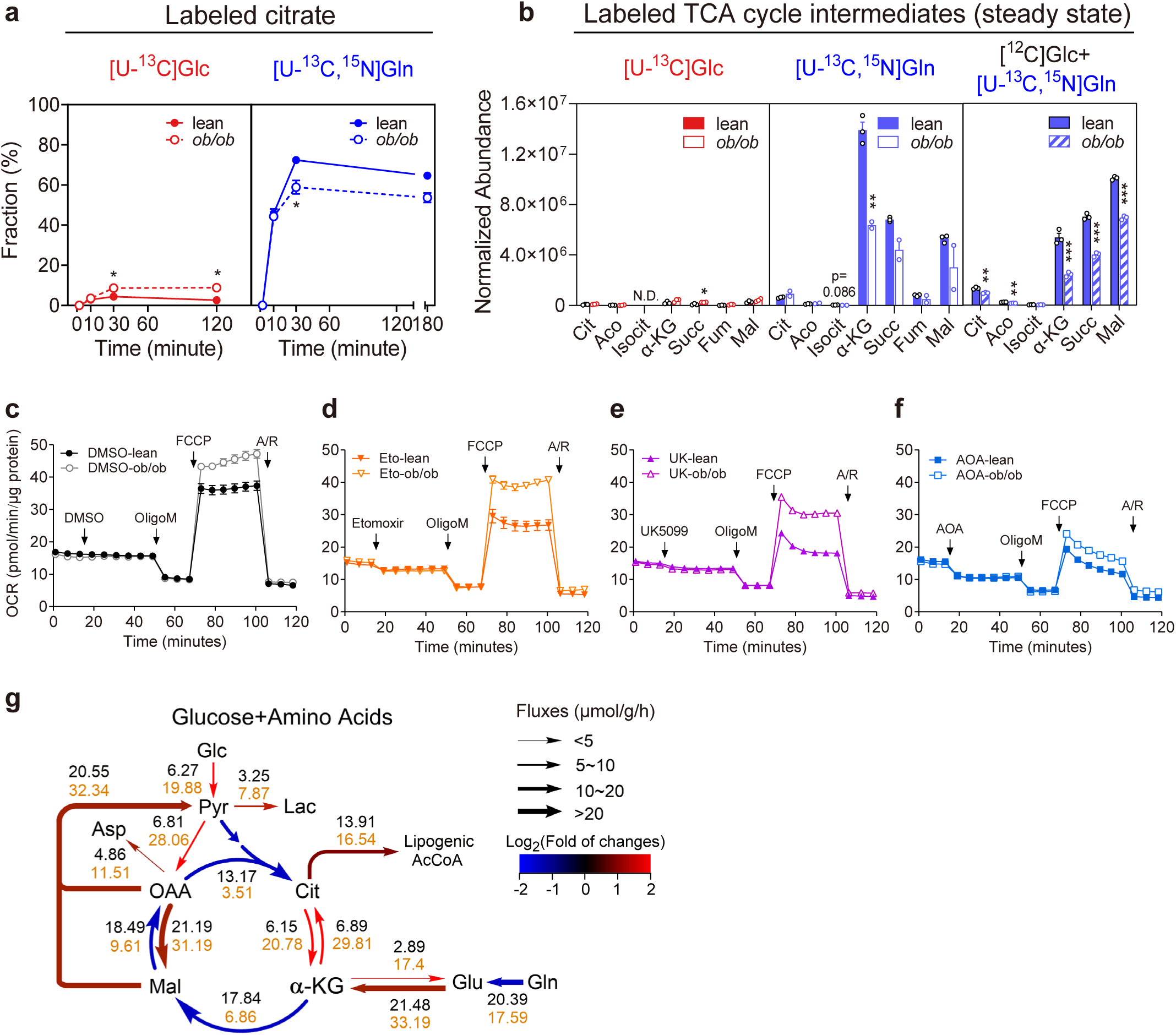
Related to Figure 3; Substrate level contribution of glucose and amino acids toward TCA cycle a, Accumulation kinetics of ^13^C isotope-labeled succinate in lean and obese primary hepatocytes traced with either 10 mM [U-^13^C]Glc or 2.5 mM [U-^13^C, ^15^N]Gln. **b,** Abundance of all ^13^C isotope-labeled TCA intermediates in primary hepatocytes at steady state. Peak areas were normalized to protein levels. **c-f,** Measurement of mitochondria respiration in primary hepatocytes, the same assay with Fig. 3c-d. Mitochondria fuel test inhibitors and mitochondria stress test compounds were used at the following concentrations: DMSO (0.1%), etomoxir (40 μM), UK5099 (40 μM), AOA (40 mM); oligomycin (OligoM, 4μM), FCCP (0.5 μM), antimycin A/rotenone (A/R, 2 μM). **g,** Estimation of TCA cycle flux based on glutamine tracing results (summary shown in Figure 3e). Best fit fluxes of primary hepatocytes isolated from lean (black) and *ob/ob* (orange) mice are presented along with each reaction. Flux rate are expressed as μmol/g cells per hour. See details in Supplementary Table 1. Data are presented as means ± SEM, outliers beyond 1.5×SD are removed; text labeled 0.05 < p < 0.1, *p < 0.05, **p < 0.01, ***p < 0.001, two-tailed unpaired Student’s *t*-test (*ob/ob vs.* lean). N.D., not detected.

**Supplementary Figure 5.**
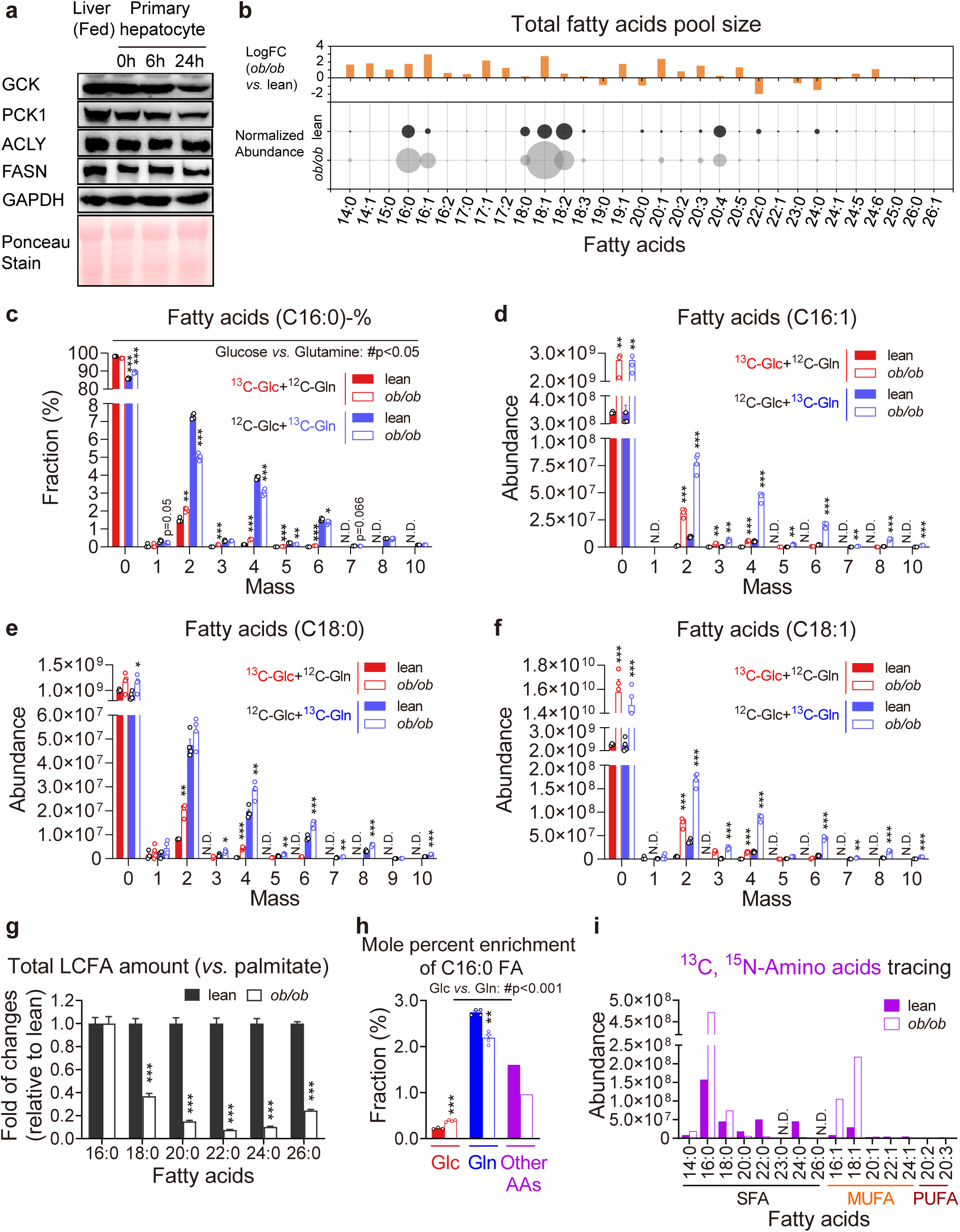
Related to Figure 4; Carbon backbone flux from glucose and amino acids toward fatty acids. **a,** Pool sizes of different fatty acids (lower panel) normalized to palmitate levels of lean mice and the log2 fold of changes (upper panel) between lean and obese primary hepatocytes. **b,** Immunoblot measurement of key enzyme expression levels among liver and isolated primary hepatocytes from the same fed mouse. 0, 6, and 24h denote the time upon the initiation of tracing studies. **c,** Fractional abundance of triglyceride palmitate (C16:0 FA) isotopomers in lean and *ob/ob* primary hepatocytes traced with either 10 mM [U-^13^C]Glc or 2.5 mM [U-^13^C, ^15^N]Gln calculated from Fig. 4b. **d-f,** Abundance of the most enriched triglyceride FA (C16:1, C18:0, C18:1) isotopomers in lean and *ob/ob* primary hepatocytes traced with either 10 mM [U-^13^C]Glc or 2.5 mM [U-^13^C, ^15^N]Gln. **g,** Fractional abundance of individual triglyceride fatty acid species in lean and *ob/ob* primary hepatocytes normalized to palmitate. Gln) by two-tailed unpaired Student’s *t*-test. N.D., not detected. **h,** ^13^C Mole percent enrichment (MPE) of palmitate (C16:0 FA) labeled by each tracer in lean mice primary hepatocytes. **i,** Combined abundance of ^13^C-labeled triglyceride fatty acid species synthesized in lean and *ob/ob* primary hepatocytes traced with [U-^13^C, ^15^N]amino acids mixture (without glutamine). SFA, saturated fatty acids; MUFA, monounsaturated fatty acids; PUFA, polyunsaturated fatty acids. Peaks areas are normalized to protein levels. Data are presented as means ± SEM, n=4 except that one outlier was discarded in the ^13^C Glc-labeled lean group; text labeled 0.05 < p < 0.1, *p < 0.05, **p < 0.01, ***p < 0.001 (*ob/ob vs.* lean) and #p<0.05 or text labeled (Glc *vs.* Gln) by two-tailed unpaired Student’s *t*-test. N.D., not detected.

**Supplementary Figure 6.**
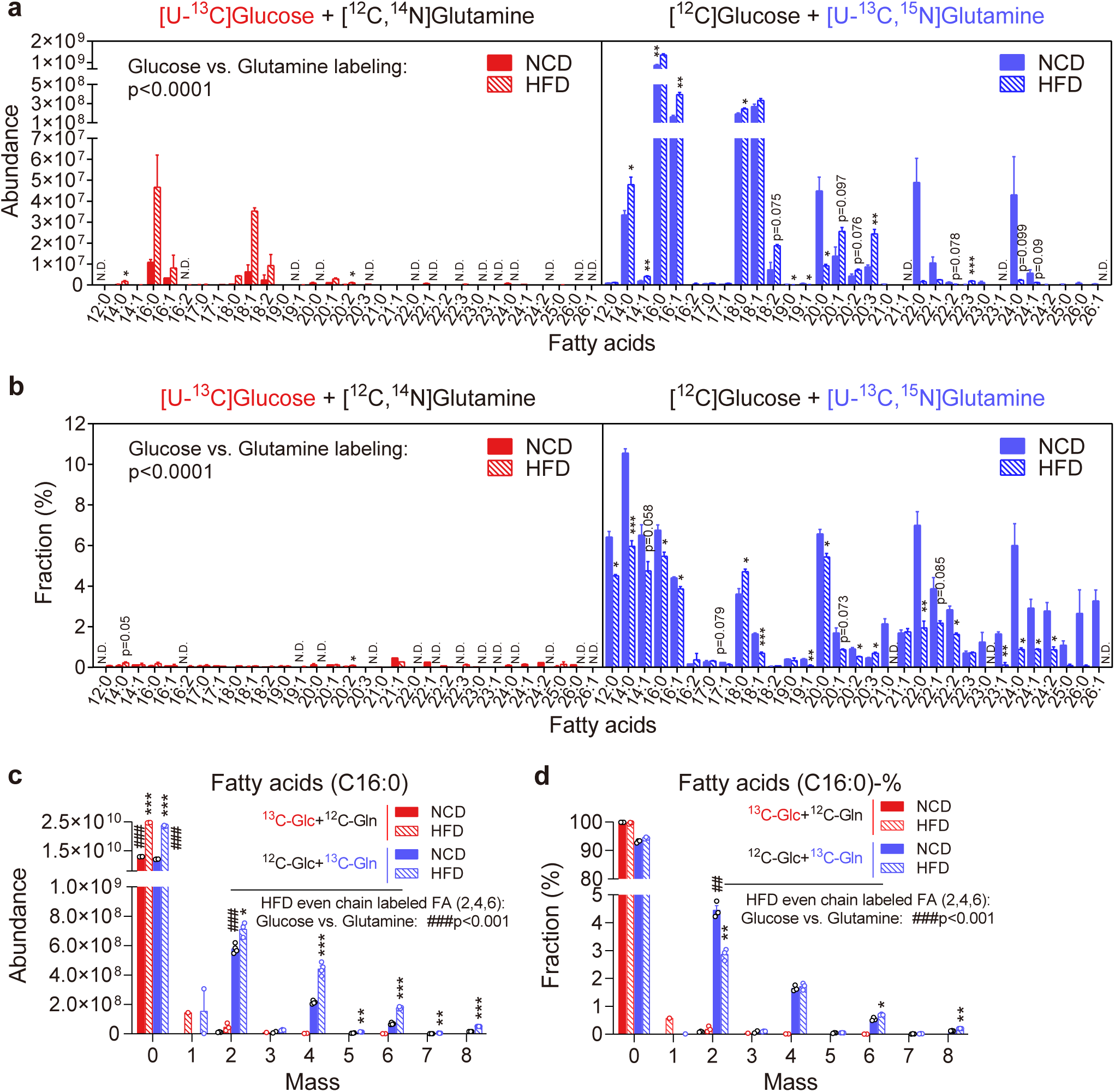
Related to Figure 4; Carbon backbone flux from glucose and amino acids toward fatty acids in primary hepatocytes isolated from normal chow diet (NCD) and high fat diet (HFD)-fed mice. **a-b,** Absolute (**a**, peak areas normalized to protein levels) and fractional (**b**) abundance of ^13^C-labeled triglyceride fatty acid isotopomers in primary hepatocytes isolated from NCD and HFD-fed mice traced with either 10 mM [U-^13^C]Glc or 2.5 mM [U-^13^C, ^15^N]Gln. The abundance of isotopomers with odd numbers of ^13^C was very low and excluded from all calculations. n=3 for each group. **c-d,** Absolute (**a**) and fractional (**b**) abundance of triglyceride palmitate (C16:0 FA) isotopomers in primary hepatocytes of NCD and HFD mice traced with either 10 mM [U-^13^C]Glc or 2.5 mM [U-^13^C, ^15^N]Gln. Abundance are normalized to within-group average to minimize the bias caused by extraction efficiency. All measurements are presented as scattered data point. Data are presented as means ± SEM with missing values removal; text labeled 0.05 < p < 0.1, *p < 0.05, **p < 0.01, ***p < 0.001 (HFD *vs.* NCD) and ###p<0.001 or text labeled (Glc *vs.* Gln) by two-tailed unpaired Student’s *t*-test. N.D., not detected.

**Supplementary Figure 7.**
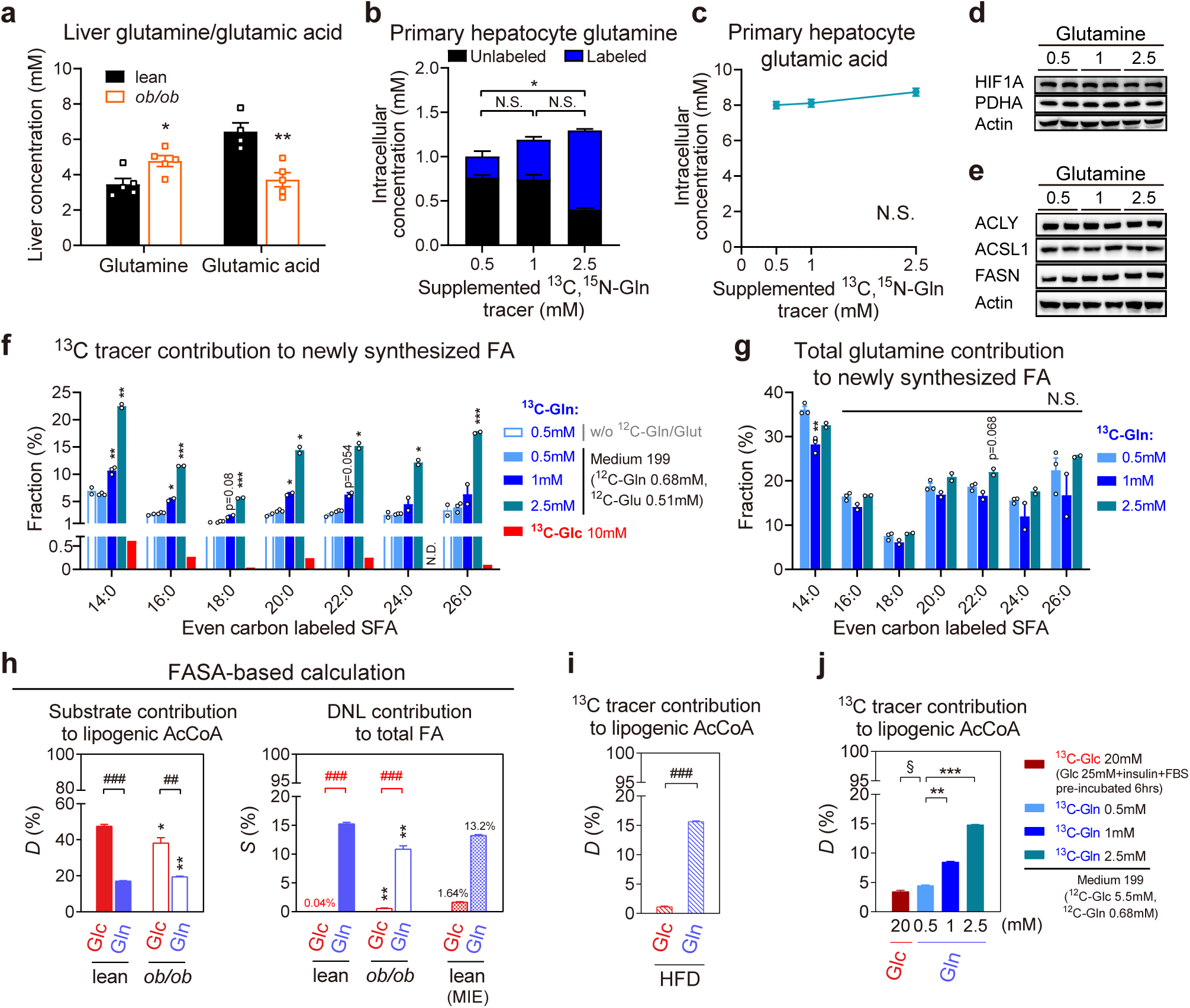
Related to Figure 4; Glutamine-DNL dose response. **a,** Quantification of glutamine and glutamic acid in liver of fed lean and *ob/ob* mice, n=5 for each group. **b,c,** Quantification of ^12^C and ^13^C glutamine (**b**), and total glutamic acid (**c**) concentration in isolated primary hepatocytes after a 6hrs labeling (steady state). n=3 for each concentration in glutamine measurement with 1 outlier in 0.5mM group removed, n=4, 3, and 3 for in glutamic acid measurement. **d,e,** Immunoblot measurement of protein levels of Hif1α and PDHα (**d**) and lipogenic enzymes (**e**) among primary hepatocyte samples traced (6hrs) with different concentrations of glutamine in complete medium (M199). **f,** Fractional abundance of saturated fatty acid (SFA) with even number of ^13^C in primary hepatocytes traced with either 0.5, 1, 2.5 mM [U-^13^C, ^15^N]Gln or 10mM [U-^13^C]Glc (single sample as a reference). n=2 for DMEM (without glutamine/glutamic acid) and n=3 for M199 complete medium. Outlier data points caused by poor fatty acid extraction efficiency were excluded. **g,** Total glutamine contribution to FA synthesis, calculated by normalizing fractional abundance of SFA in (**f**) to the [U-^13^C, ^15^N]glutamine fraction of total intracellular glutamine pool (**b**). **h,** Fractional contribution of ^13^C-labeled acetyl-CoA (AcCoA) (*D*%) toward the total lipogenic AcCoA pool, and the contribution of *de novo* synthesized fatty acid (*S*%) during the period of tracing studies toward total fatty acid pool. *D*% and *S*% are are calculated by best fitting based tool FASA (Fatty Acid Source Analysis). Red line and # text labeling indicates the estimation bias of *S*%. Lean group mass isotopomer enrichment (MIE, fractional abundances of palmitate isotopomers with even number ^13^C) serve as the lower limit reference of *de novo* lipogenesis contribution. Please refer to Figure 4h-i for comparison. **i**, *D*% of HFD primary hepatocytes (shown in Supplementary Figure 5) traced with [U-^13^C]Glc and [U-^13^C, ^15^N]Gln calculated by isotopomer aggregation method using isotopomer and even number ^13^C mole percent enrichment (MPE) of palmitate as the input. **j**, *D*% of primary hepatocytes traced with high glucose and different concentration of Gln tracer calculated by isotopomer aggregation method using data from (**f**) and (**i**). Data of Gln 1 mM group serve as the endogenous reference for *D*% normalization between 2 batch of cells. Data are presented as means ± SEM; text labeled 0.05 < p < 0.1, *p < 0.05, **p < 0.01, ***p < 0.001, # text labeled for Glc *vs.* Gln and § text labeled for low and high Glc comparison by are calculated by two-tailed unpaired Student’s *t*-test. N.S., not significant. N.D., not detected. Glc, glucose; Gln, glutamine; Glut, glutamic acid. **j,** Fractional contribution of ^13^C-labeled acetyl-CoA (AcCoA) (*D*%) toward the total lipogenic AcCoA pool, and the contribution of *de novo* synthesized fatty acid (*S*%) during the period of tracing studies toward total fatty acid pool. *D*% and *S*% are are calculated by best fitting based tool FASA (Fatty Acid Source Analysis). Red line and # text labeling indicates the estimation bias of *S*%. Lean group mass isotopomer enrichment (MIE, fractional abundances of palmitate isotopomers with even number ^13^C) serve as the lower limit reference of *de novo* lipogenesis contribution. Please refer to Figure 4h-i for comparison. **k**, *D*% of HFD primary hepatocytes (shown in Supplementary Figure 5) traced with [U-^13^C]Glc and [U-^13^C, ^15^N]Gln calculated by isotopomer aggregation method using isotopomer and even number ^13^C mole percent enrichment (MPE) of palmitate as the input. **l**, *D*% of primary hepatocytes traced with high glucose and different concentration of Gln tracer calculated by isotopomer aggregation method using data from (**f**) and (**i**). Data of Gln 1 mM group serve as the endogenous reference for *D*% normalization between 2 batch of cells. Data are presented as means ± SEM; text labeled 0.05 < p < 0.1, *p < 0.05, **p < 0.01, ***p < 0.001, # text labeled for Glc *vs.* Gln and § text labeled for low and high Glc comparison by are calculated by two-tailed unpaired Student’s *t*-test. N.S., not significant. N.D., not detected. Glc, glucose; Gln, glutamine; Glut, glutamic acid.

**Supplementary Figure 8.**
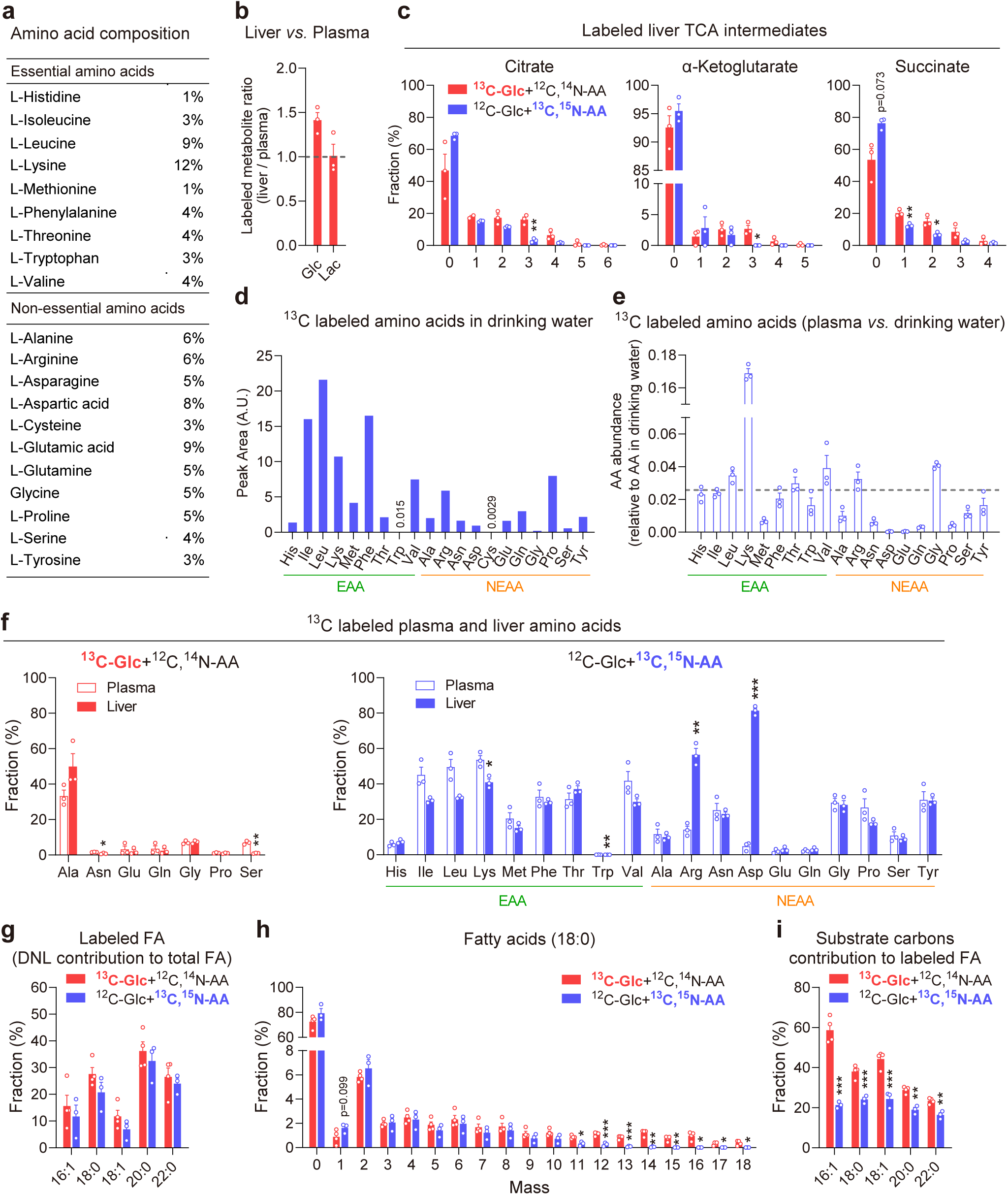
Related to Figure 5; *In vivo* metabolic flux of glucose and amino acids. **a,** Composition of [U-^13^C,^15^N]amino acid mixture used for *in vivo* isotope tracing. **b,** Ratio of ^13^C labeled glucose and lactate in the liver and plasma. **c,** Fractional abundance of ^13^C labeled TCA cycle intermediates isotopomers traced with either [U-^13^C]glucose or [U-^13^C, ^15^N]amino acids. **d,** Relative abundance of amino acids in drinking water detected by LC-MS. **e,** ^13^C labeled amino acids abundance in plasma relative to that in drinking water (**d**). **f,** Fractional abundance of ^13^C labeled amino acids in plasma (open bar) and liver (solid bar) traced with either [U-^13^C]glucose (left) or [U-^13^C, ^15^N]amino acid mixture (right). NEAA with labeling efficiency over 1% are presented. **g, h,** Total (**g**) and separative (**h**) fractional abundance of ^13^C labeled liver triglyceride fatty acids isotopomers. **i,** Fractional abundance of the labeled carbons in the newly synthesized fatty acids (**i**). Data are presented as means ± SEM; text labeled 0.05 < p < 0.1, *p < 0.05, **p < 0.01, ***p < 0.001 (Glc *vs.* AA). N.D., not detected.

**Supplementary Figure 9.**
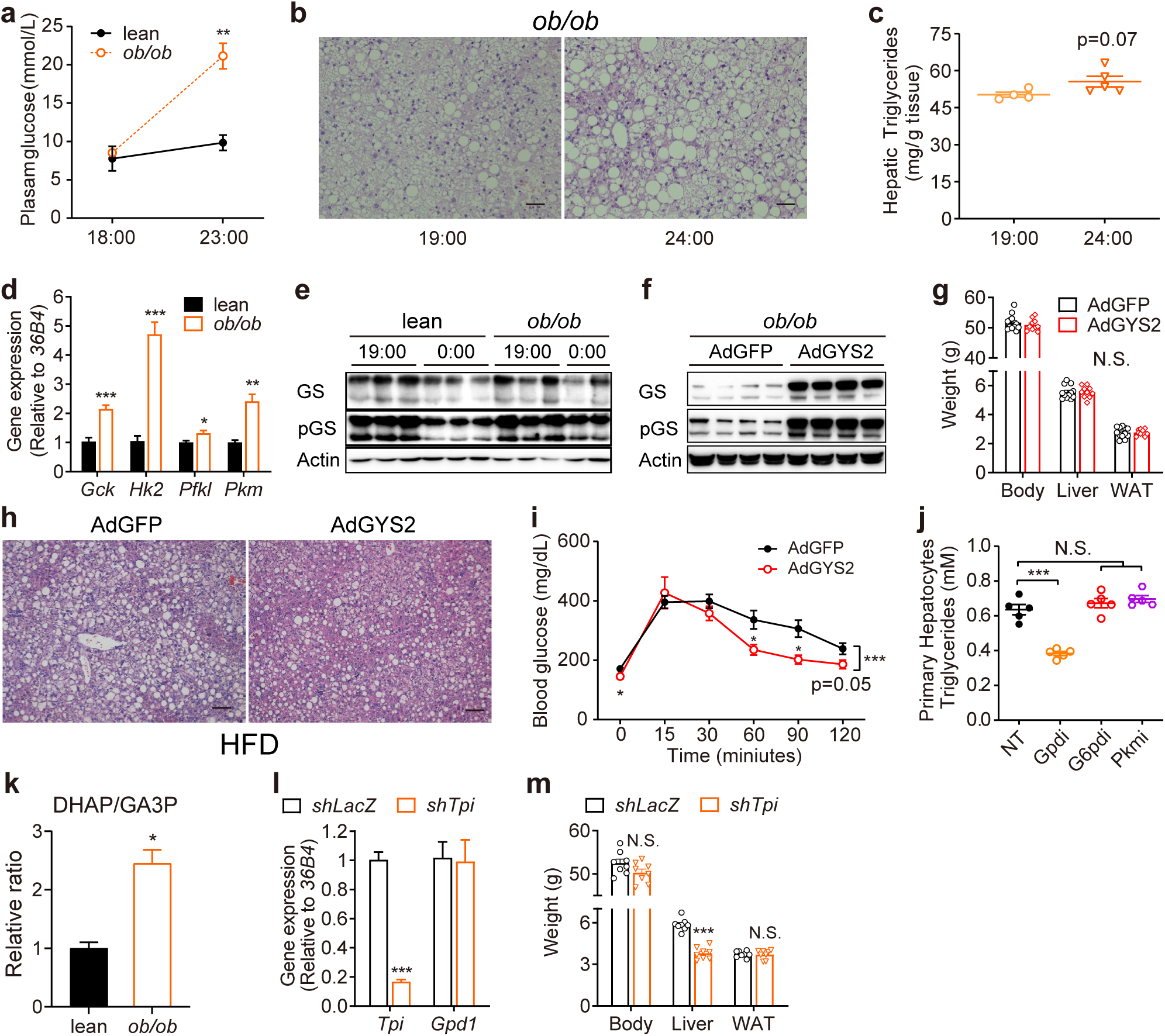
Related to Figure 6; Broad or specific inhibition of glycolytic glycerol-3P synthesis reduces hepatic triglyceride content. **a,** Measurement of changes in plasma glucose levels in lean and *ob/ob* mice in response to feeding. Lean and *ob/ob* mice were food withdrawn at ZT1 (7:00) and refed at ZT12 (18:00) for 6 hours. n=6 for the lean, and 5 for the *ob/ob* group. **b,c,** Hepatic lipid accumulation in *ob/ob* mice at ZT13 (19:00) and ZT18 (24:00) determined by H&E staining (**b**) and triglyceride measurement (**c**). n=4 and 5 for the ZT13 and ZT18 group, respectively. Scale bars, 50 μm. **d,** Measurement of glycolytic gene expression by qPCR in the liver tissue samples from refed lean and *ob/ob* mice as in (**b**), n=5. **e,** Immunoblot measurement of GYS2 protein levels and its phosphorylation state in the liver tissue samples prepared from lean and *ob/ob* mice in response to fasting and refeeding as in (**b**). **f,** GYS2 expression in primary hepatocytes prepared from *ob/ob* mice transduced with adenovirus expressing either GYS2 or control GFP. **g,** Body, liver and white adipose tissue (WAT) weight of the mice with GFP and GYS2 overexpression as described in Figure 5d-e. **h,** H&E staining of liver tissue samples from high-fat diet (HFD) mice transduced with adenoviruses expressing either GYS2 or control GFP. Tissue samples were collected 12 days post adenovirus administration. Scale bars, 50 μm. **i,** Glucose tolerance test (GTT, 2 g kg^-1^) of HFD mice with liver-specific expression of either GYS2 or control GFP seven days post adenovirus transduction. n = 7 and 6 for the GYS2 and GFP groups respectively. **j,** Triglyceride measurements of primary hepatocytes cultured in the presence or absence of inhibitors targeting GPD (ginkgolic acid 15:1, 10 μM), G6PD (6-aminonicotinamide, 1 M), or PKM (Compound 3K, 20 μM) for 24 hours, n=5. **k,** The relative abundance of DHAP over GA3P in *ob/ob* primary hepatocytes compared to the lean (calculated from flux data of Fig. 1). **l,** Measurement of *Tpi* transcript levels in *ob/ob* mice liver samples expressing shRNA targeting either *Tpi* (*shTpi*) or control (*shLacZ*), n=4. **m,** Body, liver and white adipose tissue (WAT) weight of the mice in (**l**) and Figure 6f-g. All measurements are presented as means ± SEM; *p < 0.05, **p < 0.01, ***p < 0.001, two-tailed unpaired Student’s *t-*test. N.D., not detected. N.S., not significant.

**Supplementary Figure 10.**
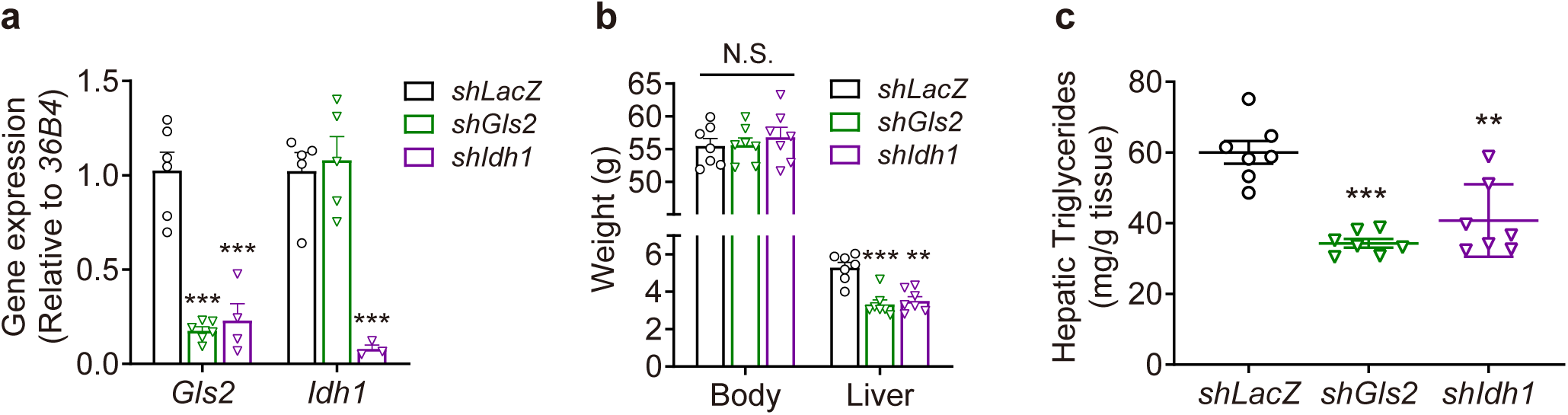
Related to Figure 7; Broad and pathway-specific manipulation of glutamine metabolism alleviates hepatic steatosis. **a,** Measurement of *Gls2* and *Idh1* transcript levels in the *ob/ob* mice with liver-specific expression of shRNA targeting *LacZ*, *Gls2* and *Idh1* mediated by adenovirus for 9 days, n=6 for sh*LacZ* and sh*Gls2*, n=4 for sh*Idh1*. **b,** Body and liver weight of liver tissues from sh*LacZ* control, sh*Gls2* and sh*Idh1 ob/ob* mice as in (**a**), n=7. **c,** Triglyceride measurement of liver tissues from sh*LacZ* control, sh*Gls2* and sh*Idh1 ob/ob* mice as in (**a**), n=7. All measurements are presented as means ± SEM; *p < 0.05, **p < 0.01, ***p < 0.001, two-tailed unpaired Student’s *t-*test. N.S., not significant. Time (minutes)

**Supplementary Figure 11.**
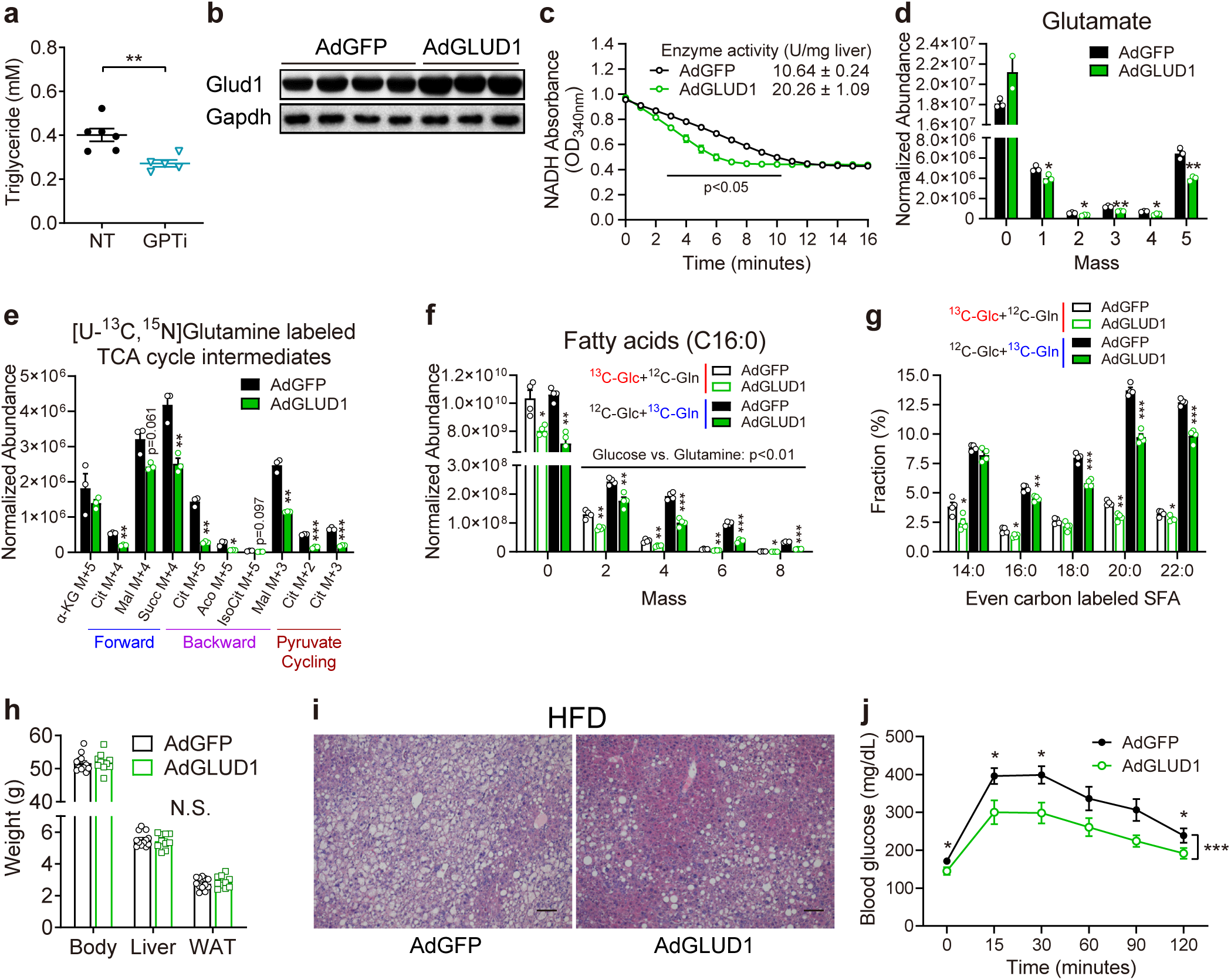
Related to Figure 7; Broad and pathway-specific manipulation of glutamine metabolism alleviates hepatic steatosis. **a,** Triglyceride measurements of primary hepatocytes cultured in the presence or absence of GPT inhibitor (3-chloro-L-alanine, 40 mM; n=5). **b,** Immunoblot measurement of GLUD1 expression in the *ob/ob* mice liver transduced with adenoviruses expressing either GLUD1 (AdGLUD1) or GFP (AdGFP). Tissue samples were collected seven days post adenovirus transduction. **c,** Measurement of GLUD1 enzyme activity from tissue samples prepared as in (**b**), n=3. **d,e,** Abundance of glutamate isotopomers (**d**) and ^13^C-labeled TCA intermediates (**e**) in [U-^13^C, ^15^N]Gln-traced (1 hour) primary hepatocytes prepared from *ob/ob* mice expressing either GFP (AdGFP, control) or GLUD1 (AdGLUD1). Primary hepatocytes were isolated three days post adenovirus transduction. Peak areas were normalized to protein levels, n=3. **f,** Abundance of ^13^C-labeled triglyceride palmitate isotopomers in either [U-^13^C]Glc or [U-^13^C, ^15^N]Gln-traced (6 hours) primary hepatocytes. Peak areas were normalized to protein levels. **g,** Fractional abundance of ^13^C-labeled triglyceride fatty acid species compared to their corresponding total pool. **h,** Body, liver and white adipose tissue (WAT) weight of the mice in (**c**) and Figure 6j-l (n=12 and 10, the AdGFP control group is shared with Supplementary Fig. 8g). **i,** H&E staining of liver sections from HFD mice expressing either GFP or GLUD1 (n=12 and 10, the control group is shared with Supplementary Fig. 8h). Scale bars, 50 μm. **j,** GTT (2 g kg^-1^ glucose) of HFD mice with liver-specific expression of GFP or GLUD1. The test was performed seven days post adenovirus administration after an overnight fasting, n=7 and 8 for the control and GLUD1 group respectively.

**Supplementary table 1.** Best fit metabolic fluxes (μmol/g cells/min) based on experimental results presented in Figure 1-3 and supplemental Figure 1-4. Flux analysis of minimal medium with multiple time points applied isotopically nonstationary ^13^C-MFA (INST-MFA) method, while that of complete medium used conventional ^13^C- MFA. Glutamate utilization by other undetected pathways were arbitrarily set as a minimal value (1.8 μmol/g cells/min). Analysis of glutamine tracing experiment in minimal medium set glycolysis influx rate as a low value (3.6 μmol/g cells/min). All the values are presented as net flux, unless specifically labeled as reverse reaction.

| Reactions | Glucose Tracing |  | Glutamine Tracing |  |  |  |
| --- | --- | --- | --- | --- | --- | --- |
|  | Minimal (Glc) |  | Minimal (Gln) |  | Complete (Glc+AA) |  |
|  | lean | ob/ob | lean | ob/ob | lean | ob/ob |
| Glycolysis (Glc→Pyr) | 10.52 | 17.57 | 3.60 | 3.60 | 6.27 | 19.88 |
| LDH (Pyr→Lac) | 1.32 | 2.82 | 4.15 | 3.38 | 3.25 | 7.87 |
| GLS (Gln→Glu) | 21.58 | 26.52 | 12.44 | 21.74 | 20.39 | 17.59 |
| GDH (Glu→α-KG) | 19.78 | 24.72 | 4.93 | 10.53 | 18.59 | 15.79 |
| GDH Reverse (α-KG→Glu) | 7.41 | 1.40 | 55.37 | 41.13 | 2.89 | 17.40 |
| PC (Pyr+CO<sub>2</sub>→OAA) | 8.33 | 5.10 | 18.37 | 10.67 | 6.81 | 28.07 |
| PDH/CS (Pyr+OAA→Cit+CO<sub>2</sub>) | 5.60 | 22.53 | 4.08 | 7.05 | 13.17 | 7.51 |
| IDH (Cit→α-KG+CO<sub>2</sub>) | -11.74 | -1.51 | -0.13 | -0.05 | -0.74 | -9.03 |
| IDH Reverse (α-KG+CO<sub>2</sub>→Cit) | 16.36 | 22.04 | 8.06 | 12.22 | 6.89 | 29.81 |
| AKGDH/SDH/FUS (α-KG→Mal+CO<sub>2</sub>) | 8.04 | 23.21 | 4.80 | 10.48 | 17.85 | 6.76 |
| MDH (Mal→OAA) | 3.31 | 10.32 | -14.29 | -3.62 | -2.70 | -25.58 |
| MDH Reverse (OAA→Mal) | 16.98 | 15.98 | 32.02 | 51.54 | 21.19 | 35.19 |
| ME/PEPCK (Mal→Pyr+CO<sub>2</sub>) | 4.73 | 12.88 | 23.30 | 21.21 | 20.55 | 32.34 |
| ACLY (Cit→AcCoA+OAA) | 17.34 | 24.04 | 4.21 | 7.10 | 13.92 | 16.54 |

## Notes

### Competing Interest Statement

The authors have declared no competing interest.

